# The dynamics and strategy of RNA replication in astroviruses

**DOI:** 10.64898/2026.01.19.700307

**Authors:** David Noyvert, Imran M. Darr, Ksenia Fominykh, Jacqueline Hankinson, Nina Lukhovitskaya, Andrew E. Firth, Valeria Lulla

**Author notes:** Correspondence: Andrew E. Firth,; Valeria Lulla.

## Abstract

Astroviruses are positive-sense single-stranded RNA viruses that cause significant disease across avian and mammalian hosts, yet their replication mechanisms remain poorly understood. The replication of astrovirus RNA occurs via a double-stranded RNA intermediate that is used as a template for the synthesis of new positive-sense RNA, which is covalently linked to the virus-encoded protein VPg. These viruses also produce a capsid-encoding subgenomic (sg) RNA that is 3′-coterminal with the genomic RNA. The mechanisms by which the astrovirus sgRNA is produced and regulated during infection have not yet been characterized. Using high throughput sequencing of RNA from cells infected with each of five different astrovirus strains, we demonstrate that the presence of a (−)sgRNA is a conserved feature of infection, supporting a premature termination model of subgenomic RNA production. A pronounced pile-up in the mapping positions of the 3ʹ ends of negative-sense RNA reads marks the precise 3ʹ terminus of the (−)sgRNA. We investigate the relative abundance and dynamics of positive and negative RNA species during virus replication and virion packaging, and perform a mutational analysis of conserved residues in the genomic and subgenomic 5ʹ termini. Together, this work elucidates the dynamics of genomic and subgenomic RNA synthesis during astrovirus infection.

**Graphical abstract:** 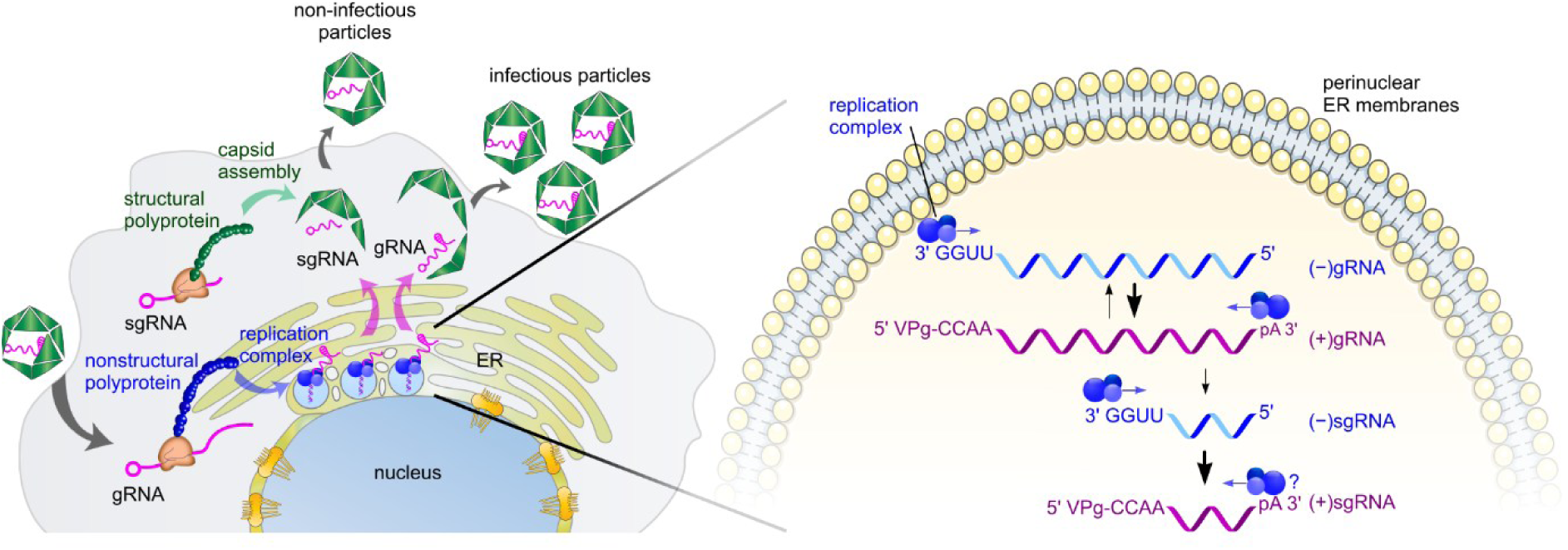

## Introduction

Human astroviruses (HAstVs) represent the least studied group of major enteric viruses ^1^. Currently, three clades of human-infecting astroviruses are recognized: classical HAstV (serotypes 1–8), and two emerging genogroups – HAstV-MLB and HAstV-VA/HMO ^2^. Ongoing surveillance studies continue to identify novel astroviruses in many mammalian species, indicating the potential for future zoonoses ^3^. In Asia, Africa and South America, astroviruses represent an overlooked cause of diarrhea in children, reaching 35% of enteric infections ^4^. Non-classical MLB and VA astroviruses often present extra-intestinal pathogenicity, resulting in meningitis and encephalitis in the immunocompromised and elderly ^1,5–8^. Astrovirus infections are also associated with prolonged virus shedding and diarrhea in immunocompromised children ^9^. Despite their significant impact on public health, no vaccines or antiviral agents have been approved to treat astrovirus infection.

Astroviruses are non-enveloped positive-sense single-stranded RNA viruses, whose genomes are polyadenylated at the 3ʹ end and covalently linked to a viral protein, VPg, at the 5ʹ end. The genome contains a 5ʹ untranslated region (UTR), a 3ʹ UTR, and three or four protein-coding open reading frames (ORFs). ORF1a and ORF1b encode the viral replication polyproteins and contain domains corresponding to N-terminal domain ^10^, transmembrane protein, protease, VPg, RNA dependent RNA polymerase (RdRp), and highly variable C-terminal protein ^11–13^. Two ORFs are expressed from subgenomic (sg) RNA: ORF2 encodes the structural polyprotein (capsid) ^11^, and ORFX (present in many species) encodes XP, a viroporin ^14^.

Synthesis of sgRNAs is a common strategy used by positive-sense RNA viruses to express 3ʹ-proximal genes ^15^. Viral sgRNAs often encode proteins needed for virus particle assembly, release and pathogenesis ^16^. Expression of sgRNAs usually depends on sg promoter elements, often comprising *cis*-acting RNA elements that can be located both upstream and downstream of the transcription start site ^17^.

Astrovirus RNA replication is poorly understood and mostly inferred from other positive-sense RNA viruses. Positive-sense RNA viruses employ three general strategies for sgRNA production. The first one is utilized by members of the *Togaviridae* and *Bromoviridae* families and is based on internal initiation ^15^: full-length (−)gRNA intermediate is used as a template for synthesis of both full-length (+)gRNA and the shorter (+)sgRNA. Here, synthesis of sgRNA depends on an internal promoter located within the (−)RNA (Fig. 1A). The second strategy is common in the *Tombusviridae* and *Nodaviridae* families ^18–20^. Here, premature termination can occur during (−)RNA synthesis, leading to generation of a subgenomic-sized negative-sense RNA, (−)sgRNA, that is used as the template for (+)sgRNA synthesis (Fig. 1B). The third option, discontinuous transcription, is used in the *Coronaviridae* and *Arteriviridae* families. This unusual strategy results in the production of a nested set of 3ʹ-coterminal sgRNAs that also contain a conserved 5ʹ leader identical in sequence to the 5ʹ end of the genome ^21,22^. It was previously proposed that astroviruses use the first mechanism to produce sgRNA; however, this had not been experimentally confirmed ^23,24^.

**Figure 1.**
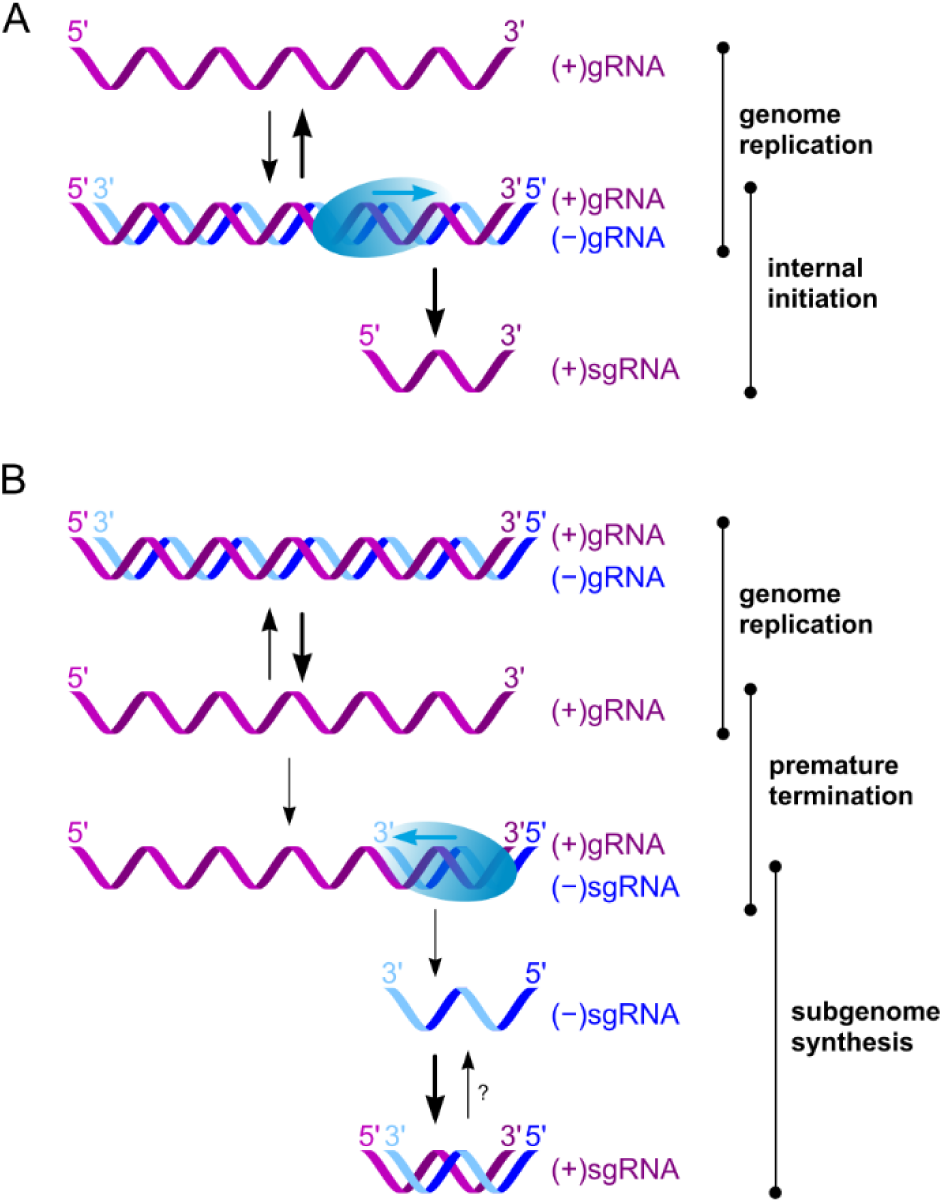
Schematic depicting RNA replication strategies in positive-sense RNA viruses. (**A**) The genomic positive-sense RNA, (+)gRNA, serves as a template to produce negative-sense genomic RNA, (−)gRNA, and the latter serves as a template for both (+)gRNA and (+)sgRNA synthesis. (**B**) The (+)gRNA serves as a template for (−)gRNA synthesis, and a negative-sense sgRNA, (−)sgRNA, is also produced via premature termination. (−)sgRNA is used as a template for (+)sgRNA synthesis. In some cases, it is possible that (+)sgRNAs are further amplified, i.e., used as templates for new (−)sgRNA synthesis.

We previously created replicons for HAstV1, MLB1 and MLB2 astroviruses that rely on minimal RNA elements required to produce and translate subgenomic messages ^14,25^. Surprisingly, these RNA elements appear to be quite long and span 46–150 nucleotides downstream of the ORF2 initiation site. In addition, synonymous site conservation analyses showed substantial conservation – indicative of functionally important elements – extending for ∼170 nt upstream of the sgRNA initiation site ^14^. Together, these observations suggest that sgRNA synthesis in astroviruses may be regulated by RNA-RNA and/or RNA-protein interactions located at both sides of the transcription termination/initiation site.

Here, we overturn the previously assumed model of astrovirus sgRNA synthesis. We identify a (−)sgRNA species ubiquitously present during astrovirus replication (characteristic of premature termination sgRNA synthesis), we analyze terminal features of RNA species, and we evaluate the dynamics of gRNA/sgRNA production during astrovirus infection.

## Results

### Identification of a subgenome-sized negative-sense RNA species

Two distinct fundamental mechanisms to produce viral sgRNAs are internal initiation and premature termination (Fig. 1). These can be distinguished by the absence or presence, respectively, of a (−)sgRNA intermediate. Astrovirus infection is characterized by the presence of two major RNA species detectable by Northern blotting (Fig. 2A): (+)gRNA (∼6.8 kb) and (+)sgRNA (∼2.5 kb). However, detection and quantification of less abundant RNA species, particularly at earlier replication stages, require more sensitive methods with a higher dynamic range.

**Figure 2.**
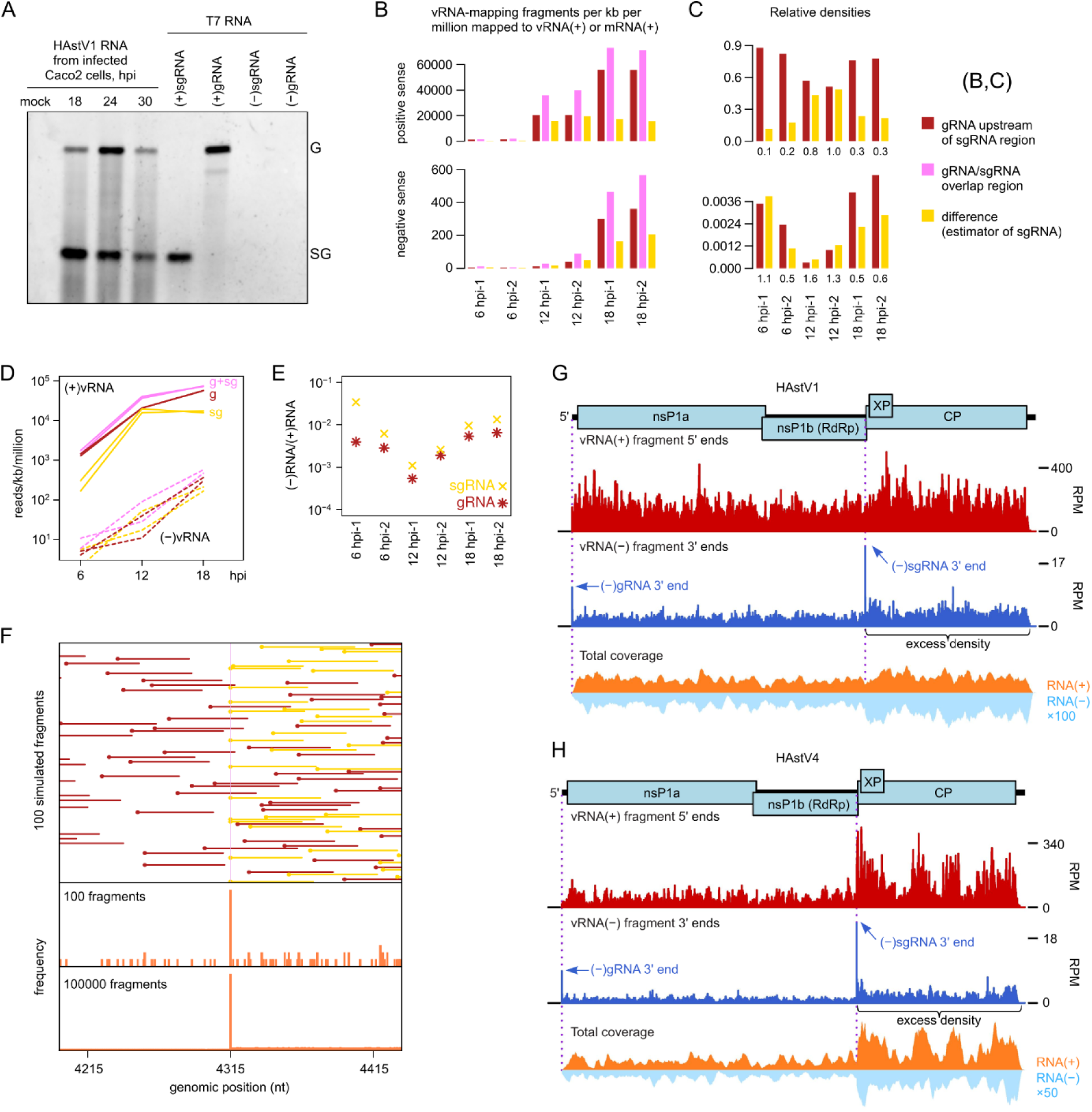
RNA-seq analysis of astrovirus-infected cells. (**A**) Northern blot analysis of poly(A)-selected RNA isolated from infected Caco-2 cells (500 ng per sample); 100 ng of genomic and 25 ng of subgenomic *in vitro* transcribed T7 HAstV1 RNA were used as a control. (**B**) Caco-2 cells were infected with HAstV1 at MOI 5 and harvested at 6, 12 or 18 hpi in duplicate. Bar graphs show the density of mapped fragments in the gRNA/sgRNA-overlap region (pink), upstream of the sgRNA region (red), and the difference (yellow). Coverage is quantified as fragments per kilobase per million fragments mapped to vRNA(+) or host mRNA(+). Fragments mapping to the sgRNA region may derive from either gRNA or sgRNA; the difference (yellow) in density between the sgRNA and non-sgRNA regions was used to estimate the relative abundance of sgRNA, whereas the density in the non-sgRNA region (red) was used to estimate the relative abundance of gRNA. **(C)** Relative densities of (+)gRNA, (+)sgRNA, (−)gRNA and (−)sgRNA. Numbers below bars show the estimated sgRNA:gRNA ratio (1 d.p.). (**D**) Same data as panel (B), plotted on a log scale. (**E**) Estimated (−):(+) ratio for gRNA and sgRNA species. (**F**) Length 6785 nt gRNAs (red), together with sgRNAs (yellow) beginning at nt 4315, in a 1:1 ratio, were randomly fragmented *in silico*, using a fixed 1/60 probability of cleavage between any adjacent pair of nucleotides. Then fragments in the 50–70 nt length range were selected. The top panel schematically illustrates the first 100 fragments that map at least partly within the region from 120 nt 5ʹ to 120 nt 3ʹ of nt 4315. The middle panel shows a histogram of the 5ʹ end positions of these 100 fragments. The bottom panel shows an equivalent histogram for the first 100,000 similarly selected fragments. (**G**) Caco-2 cells were infected with HAstV1 at MOI 5 and harvested at 18 hpi, with proteinase K (PK) treatment. Histograms show positions of 5ʹ ends of fragments mapping to vRNA(+) (red), 3ʹ ends of fragments mapping to vRNA(−) (blue), and total coverage of vRNA(+) and vRNA(−) (orange and pale blue, respectively). For the 5ʹ/3ʹ-end histograms, counts were normalized to fragments per million fragments mapped to vRNA(+) or host mRNA(+) (’reads per million’; RPM). For the 3ʹ end plot, the histogram shows the 3ʹ ends of negative-sense fragments, corresponding to 5ʹ ends of the positive-sense reverse complements of the fragments. For the total coverage plot, the *y*-axis scale is arbitrary, but vRNA(−) coverage is scaled relative to vRNA(+) coverage by the indicated factor to aid visualization. The figure is for tech. rep. 1, 18 hpi, PK, biol. rep. 2; see Figs S5-S6 and S8-S11 for 5ʹ/3ʹ-end and total coverage histograms for all 16 samples. **(H)** As for panel (G) but for HAstV4, 24 hpi, biol. rep. 1; see Figs S13-21 for the full set of histograms for HAstV4, MLB1, MLB2 and VA1 infections.

To evaluate which mechanism astroviruses utilize, we performed high throughput sequencing of RNA (RNA-seq) extracted from astrovirus-infected cells. Initially, we analyzed HAstV1 infection at 6, 12, and 18 hours postinfection (hpi). The total coverage of RNA-seq reads on the HAstV1 (+) and (−) strands is shown in Fig. S1. A noticeable excess density in the sgRNA region for negative-sense reads strongly indicates the presence of a (−)sgRNA. Mapped RNA-seq density often varies erratically along transcripts as a result of biases during library preparation, including ligation and PCR biases. Therefore, we also calculated the mean RNA-seq densities in the sgRNA region and in the region of gRNA not overlapped by sgRNA. We used the difference in density between the 3ʹ sgRNA/gRNA-overlap region and the 5ʹ gRNA-only region as an estimator of the sgRNA density (Fig. 2B-D).

The increase in (+)gRNA relative to (+)sgRNA from 12 hpi to 18 hpi (Fig. 2C) is understandable, as we might expect elevated production of gRNAs for packaging at late timepoints. The parallel increase in (−)gRNA relative to (−)sgRNA from 12 hpi to 18 hpi may reflect increased production of (−)gRNA to drive the increased production of (+)gRNA. At early timepoints, we might expect RNA synthesis to initially favor gRNA replication to drive production of replication complexes, followed later by sgRNA synthesis to drive capsid protein production and virion formation. One interpretation of Fig. 2C is that at 6 hpi (−)sgRNA synthesis is more advanced than (+)sgRNA synthesis, supporting the second sgRNA synthesis strategy (Fig. 1B). However, caution is needed because another possible explanation for the low (+)sgRNA:(+)gRNA ratio at 6 hpi may be potential leftover input (+)gRNA from the infection that is not being efficiently replicated to (−)gRNA. In addition, at high MOI infection, the proportion of defective RNA species is usually higher ^26^, which can also influence both (+) and (−) RNA ratios. Moreover, as a proportion of total virus (+)RNA plus host mRNA, (−)sgRNA density increases substantially from 12 hpi to 18 hpi during which (+)sgRNA slightly decreases, suggesting late timepoint decoupling between (−)sgRNA and (+)sgRNA synthesis, or even that new (−)sgRNAs may be synthesised from (+)sgRNA templates (Fig. 2D). We also looked at (−)RNA:(+)RNA ratios (Fig. 2E). Although the (−)sgRNA:(+)sgRNA was higher at 6 hpi than at 12 hpi, it was still only 0.034/0.006 for the two repeats, i.e., even at 6 hpi there appeared to be far more (+)sgRNA than (−)sgRNA (but see below). Thus, we cannot say from these data whether the observed (−)sgRNA is a precursor to (+)sgRNA or whether (−)sgRNA is produced from a (+)sgRNA precursor (either accidentally or for sgRNA amplification), or perhaps both occur at different stages of infection.

Besides mapping total RNA-seq coverage across the virus (+) and (−) strands, it is also possible to assess the positions of the 5ʹ and/or 3ʹ ends of mapped reads. This can provide supporting evidence for the start and stop sites of transcripts because, following random fragmentation, reads are more likely to end at a pre-fragmentation transcript end than at any given random fragmentation point, leading to a single-nucleotide peak in a histogram of 5ʹ or 3ʹ read-end mapping positions (Fig. 2F) ^27^. Since the 5ʹ ends of astrovirus gRNAs (and likely also sgRNAs) are covalently linked to the viral protein VPg ^28^, these transcript termini are probably unavailable for RNA-seq adaptor ligation (even after protease treatment). Furthermore, the 3ʹ ends of these transcripts are polyadenylated and of variable length, precluding a 3ʹ end pile-up. As expected, no peaks were observed at the 5ʹ end of gRNA or sgRNA when the 5ʹ ends of positive-sense HAstV1 reads were mapped to the viral genome (Fig. S2). In contrast, the 3ʹ ends of (−)gRNA and (−)sgRNA species (corresponding in position to the 5ʹ ends of (+)gRNA and (+)sgRNA, respectively) are expected to be available for adaptor ligation. Nonetheless, histograms of the mapping positions of the 3ʹ ends of negative-sense HAstV1 reads were inconclusive (Fig. S3): a modest peak (the highest peak genome-wide but not greatly above background) could be observed at the sgRNA start site for the two 18 hpi repeats but not at earlier timepoints (likely due to too few negative-sense reads; Table S1).

Therefore, we repeated the analysis, this time at 18 and 24 hpi timepoints. In addition, we prepared samples with or without proteinase K (PK) treatment. The initial idea was to test whether this would completely remove VPg to allow adaptor ligation to transcript 5ʹ termini. Unfortunately, use of PK did not yield clear peaks at the 5ʹ ends of (+)gRNA or (+)sgRNA in histograms of read 5ʹ-end mapping positions (Figs S4 and S5), likely due to incomplete removal of VPg. Unexpectedly, PK treatment resulted in a substantial increase in the ratio of (−)RNA to (+)RNA reads (Fig. S6), suggesting that this treatment may increase the release of (−)RNA from replication complexes.

With these new datasets, we again saw excess read density in the sgRNA region for (−)RNA for all 16 samples (2 time points, with or without PK, 2 biological repeats, 2 technical repeats) (Figs S6-S8). For (+)RNA, the excess density in the sgRNA region was less pronounced at 18 hpi and not apparent at 24 hpi, likely due to (+)RNA at the latter timepoint being dominated by (+)gRNA destined for packaging or already packaged into virus particles. PK treatment increased the (−)RNA:(+)RNA ratio by 4.4 to 12.5 fold (mean 8.2) (Table S2). Even accounting for the mean 8.2 increase upon PK treatment, the 0.006–0.034 (−)sgRNA:(+)sgRNA ratio observed at 6 hpi (see above; Fig. 2E) would only correct to 0.05–0.28 (i.e. still less (−)sgRNA than (+)sgRNA at 6 hpi). With PK treatment, the (+)gRNA/(−)gRNA ratio was in the range 48–118 (mean 78 and 110 at 18 and 24 hpi, respectively), and the (+)sgRNA/(−)sgRNA ratio was in the range 8–34 (mean 20) at 18 hpi, whereas at 24 hpi it could not be determined due to insufficient RNA-seq density difference between the gRNA/sgRNA overlap region and the gRNA-only region (Figs S6-S8).

In the histograms of read-end mapping positions, we observed a clear peak at the 3ʹ end of (−)gRNA and (−)sgRNA (i.e. corresponding in position to the 5ʹ end of (+)gRNA and (+)sgRNA) for some but not all samples (Figs S9-S10). Notably, this was particularly clear in the 18 hpi PK-treated repeat 2 sample (Fig. 2G).

Next, we extended the RNA-seq analysis to another classical human astrovirus strain, HAstV4, besides newly emerged astrovirus strains VA1, MLB1 and MLB2 (Fig. 2H; Figs S11-S20). All infections were performed with an MOI of 5, cells were harvested at 24 hpi, and we used the PK treatment for all samples. In all cases, there was a clear excess density in the sgRNA region both for negative-sense and positive-sense reads (Fig. S11). The 3ʹ ends of the (−)gRNA and (−)sgRNA were particularly clear in the HAstV4 histograms (Fig. 2H; Figs S17). For VA1, a clear peak was observed at the 3ʹ end of the (−)gRNA, but not the (−)sgRNA (Fig. S20), potentially reflecting a combination of the low number of mapped (−)RNA reads (Table S3) and the infection stage. Similarly, for MLB1 and MLB2, the low number of mapped (−)RNA reads precluded a useful read-end mapping analysis (Figs S18-S19, Table S3). The low number of mapped (−)RNA reads for MLB1, MLB2 and VA1 compared to HAstV4 (∼100-fold difference; Table S3) may indicate that these viruses are in different replication stages at 24 hpi and/or that MLB1, MLB2 and VA1 grow less efficiently than HAstV1 and HAstV4 in these cell lines ^25^.

Where present (i.e., for HAstV1 and HAstV4), the 3ʹ (−)gRNA peak corresponds to the sequence 3ʹ-GGUUC and the 3ʹ (−)sgRNA peak corresponds to the sequence 3ʹ-GGUUU. We also tested all samples for the possibility of untemplated residues at the 3ʹ end of the (−)gRNA or (−)sgRNA transcripts by querying the raw RNA-seq reads (see Methods), but we found no evidence for untemplated residues. The 3ʹ (−)gRNA sequence corresponds to the known 5ʹ end of the (+)gRNA, 5ʹ-CCAAG. Notably, the 3ʹ (−)sgRNA sequence contains two additional terminal nucleotides compared to the previously determined 5ʹ end of the HAstV1 sgRNA ^29^. With our revised sgRNA 5ʹ end position, the 5ʹ ends of (+)gRNA and (+)sgRNA start with the common sequence 5ʹ-CCAA (Fig. 3A). The gRNA 5ʹ-CCAA is a common feature of our five genomes (HAstV1, HAstV4, MLB1, MLB2, VA; see Methods). Moreover, the sgRNA CCAA sequence is highly conserved across mammalian astroviruses (Fig. 3B-C), suggesting that 5ʹ-CCAA is a common feature of mamastrovirus gRNAs and sgRNAs.

**Figure 3.**
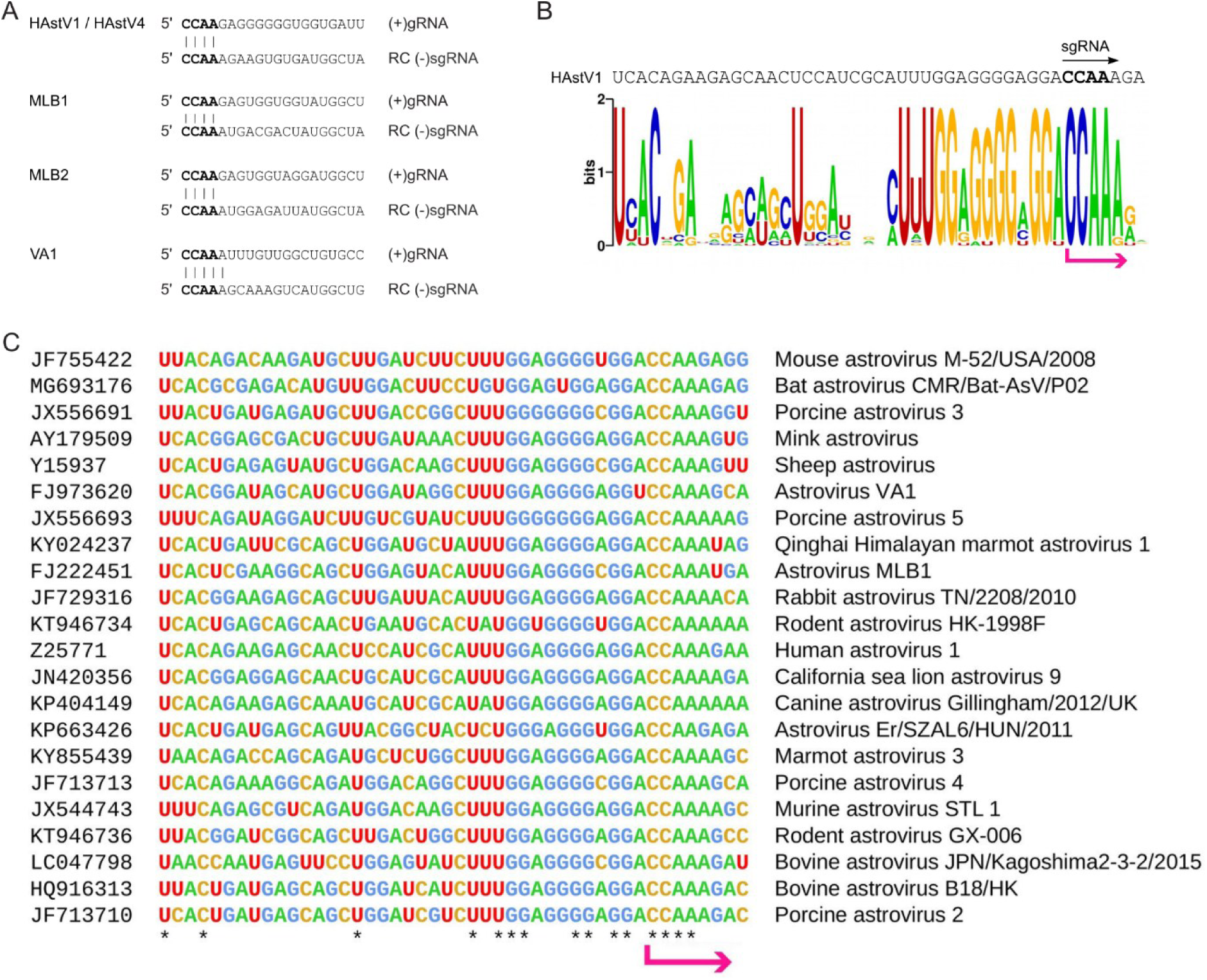
Conserved sequences at the gRNA and sgRNA start sites. (**A**) 5ʹ-terminal sequences of the (+)gRNA and inferred (+)sgRNA for the HAstV, MLB and VA astrovirus strains. (**B**) Sequence logo illustrating nucleotide conservation around the inferred 5ʹ-CCAA sgRNA start site (pink arrow), based on an alignment of 22 diverse mammalian astrovirus sequences. (**C**) Alignment of representative genus *Mamastrovirus* sequences showing the region around the sgRNA 5ʹ end (pink arrow). Asterisks indicate completely conserved positions in the alignment. See Fig. 1a of Lulla *et al.* (2020)^14^ for a phylogenetic tree.

### Functional analysis of the conserved CCAA sequence using a dual-luciferase replicon assay

To address the importance of the CCAA sequence (hereafter referred to as the CCAA motif) in both gRNA and sgRNA synthesis, we modified recently developed astrovirus subgenomic replicons for HAstV1 ^14^ and MLB2 ^25^ to include a genomic reporter (firefly luciferase) in the nsP1a region, similar to a recently published reporter (Fig. 4A) ^30^. The replication-deficient RdRp knockout control (RdRp-KO, GDD→GNN) was used in all replicon assays to evaluate baseline reporter translation levels in both replicons (Fig. 4B-E). We should note a potential limitation of the genomic reporter in that the luciferase has been inserted into a critical location, namely the N-terminal domain of nsP1a, that ensures the delivery of non-structural polyprotein to the ER membranes ^10^. This likely results in gRNA synthesis levels that are not directly comparable to those during astrovirus infection. However, due to statistically significant differences in the WT/GNN ratio, we (Figs 4B and 4D) and others ^30^ can still assess regions involved in gRNA replication.

**Figure 4.**
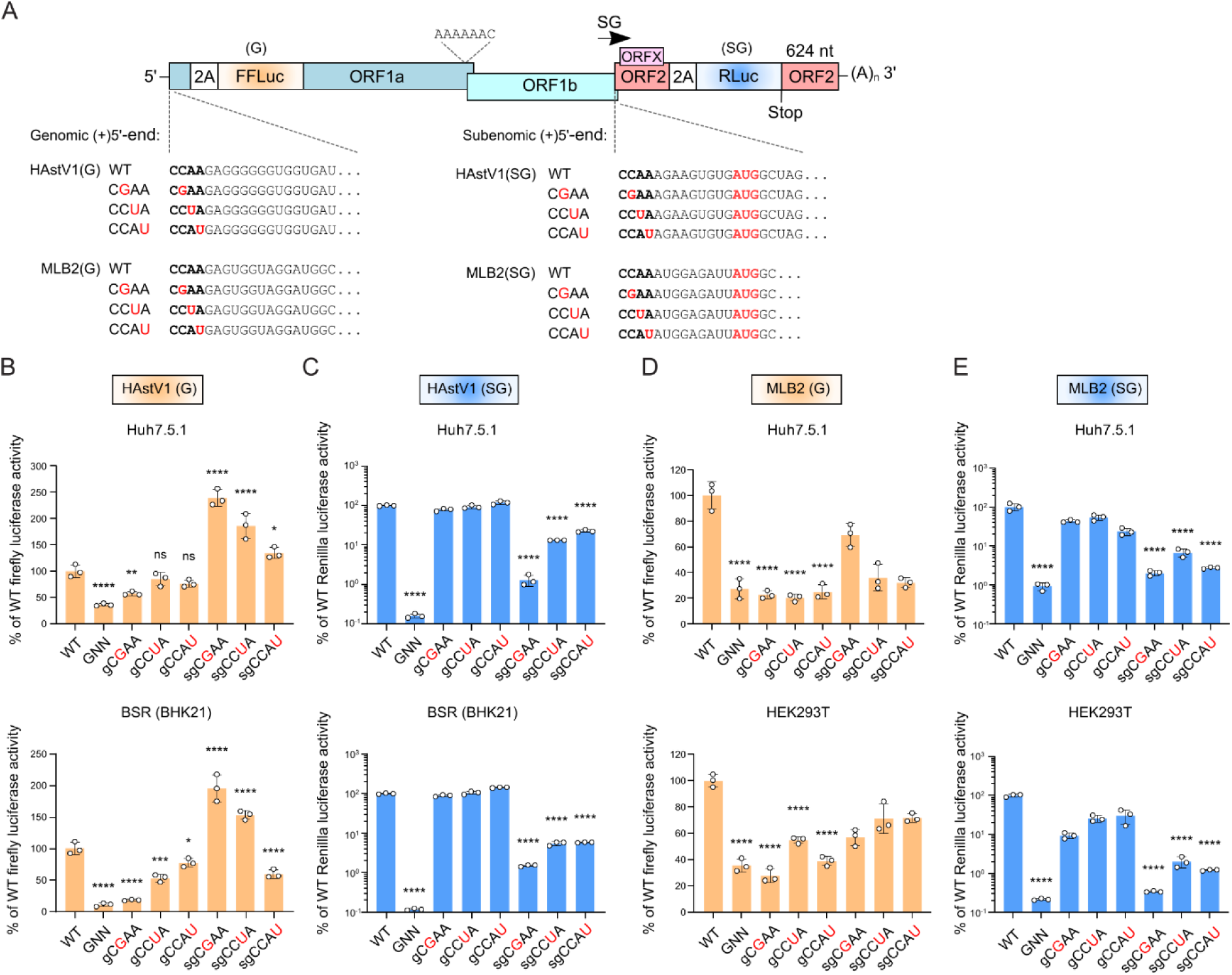
The development of the dual-luciferase replicon system and analysis of the functional importance of the conserved terminal (C)CAA sequence in genomic and subgenomic RNA production. (**A**) Schematic representation of dual-luciferase astrovirus replicons and (C)CAA mutants. (**B-E**) Relative activities for HAstV1 (B-C) and MLB2 (D-E) replicons, measured at peak signal of genomic (firefly luciferase, linear scale) and subgenomic (Renilla luciferase, log scale) reporters in the two indicated cell lines. The RdRp-KO (GDD→GNN) mutant was used as a replication-deficient control. For each mutant, both genomic and subgenomic activities were measured and compared against those of the WT replicon. Data are mean ± SD (*n* = 3 independent experiments). \**p* < 0.05, \*\**p* < 0.01, \*\*\**p* < 0.001, \*\*\*\**p* < 0.0001, ns, nonsignificant using one-way ANOVA test against WT replicon. ORF, open reading frame; FFLuc, firefly luciferase; RLuc, Renilla luciferase.

We selected the second to fourth residues from the conserved CCAA 5′-terminal g/sgRNA sequence for mutational analyses, leaving out the first C since it is likely covalently linked to VPg ^28^ and therefore cannot be assessed independently of its protein-linkage function. A comparison of (C)CAA mutants to WT and RdRp-KO control in most assays showed that they are essential for both genomic and subgenomic synthesis, with the second C nucleotide being the most impactful, followed by the third (A) and/or fourth (A) positions. These results suggest that the RdRp can use the same genomic and subgenomic RNA binding site on the 3′(−)GGUU sequence. For both HAstV1 and MLB2 replicons, we observe the same pattern in two different cell lines (Fig. 4B-E). It is also possible that (C)CAA mutations affect VPg linkage (e.g. if the 3ʹ-GGUU sequence templates the cytidylylation of VPg) in addition to the RdRp recognition defects. Interestingly, mutation of the second and third positions in C**CA**A for the HAstV1 sgRNA reporter increased expression of the gRNA reporter, suggesting competition between the gRNA and sgRNA CCAA motifs (Fig. 4B). This effect was not observed in MLB2 replicons (Fig. 4D-E), potentially due to lower replication levels ^25^.

### Analysis of the dynamics of g/sg RNA production during astrovirus infection

To further assess temporal regulation, we performed a series of analyses to evaluate and quantify the relative abundance of all four RNA species (genomic and subgenomic RNAs for both positive and negative sense) in infected cells and to quantify packaging preferences for these RNA species, using strand-specific RT-qPCR (Fig. 5A). We identified all four major RNA species in cells during high MOI infection (Fig. 5B-C) and from media-derived packaged virions in low MOI multistep growth conditions (Fig. 5D).

**Figure 5.**
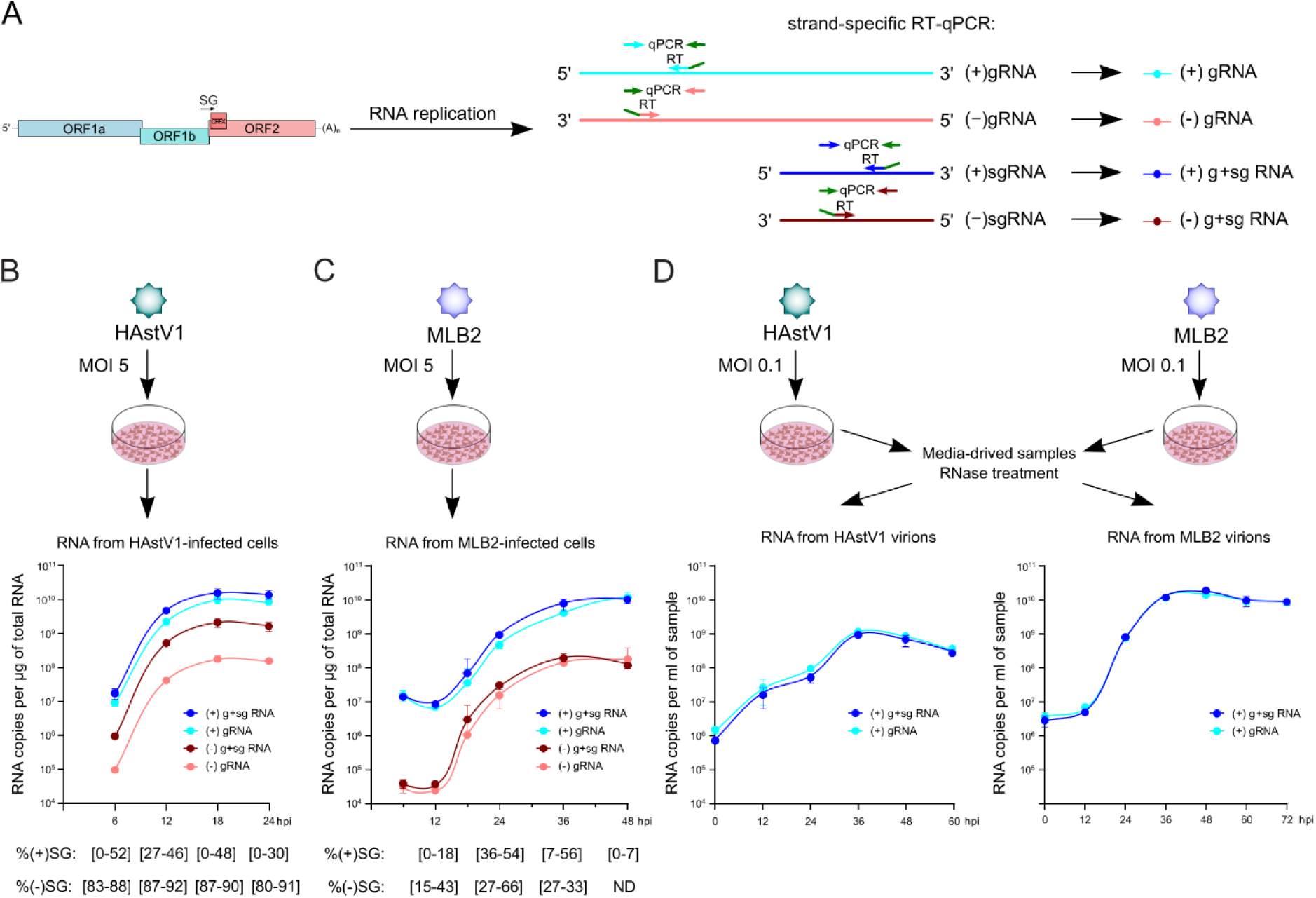
Analysis of intra- and extracellular astrovirus RNAs. (**A**) Schematic representation of strand-specific RT-qPCR. Each RT primer contains the adapter sequence (shown in green), which is subsequently used in qPCR amplification, thus reducing unspecific binding and off-target effects. (**B**) Experimental setup and RNA replication dynamics in HAstV1-infected Caco-2 cells. (**C**) Experimental setup and RNA replication dynamics in MLB2-infected Huh7.5.1 cells. (**D**) Experimental setup and RNA analysis of HAstV1 and MLB2 virions following low MOI (0.1) infection. Data are mean ± SD (*n* = 3). There is no statistically significant difference between (+)gRNA and (+)g+sgRNA samples at all time points, suggesting the absence of packaged (+)sgRNA in these conditions (e.g., low MOI, and RNase treatment). The (−)RNA was not detected in these samples.

Similar to the RNA-seq data (Fig. 2B-E), and consistent with other (+)ssRNA viruses, we found RNA replication to be asymmetric, with substantially higher levels of (+)RNA than (−)RNA (Fig. 5B-C). We observe larger differences in sg/g RNA abundance for negative-strand RNAs across all time points in HAstV1-infected cells (Fig. 5B), which were less apparent in NGS libraries (Fig. 2B-E), suggesting that differences in sample preparation and/or analysis could affect the quantification of low-abundance (−)RNA intermediates. Interestingly, this is not the case in MLB2 infection (Fig. 5C).

RNA packaging during virus release favors full-length infectious gRNA over sgRNA in both astrovirus strains (Fig. 5D), suggesting the existence of a packaging signal in the 5′UTR and/or nsP1a/b coding region. Consistent with the RNA virus replication strategy, where (−)RNA species serve as replication intermediates, only (+)RNA was detected in media-derived virus samples (Fig. 5D).

### Analysis of RNA in astrovirus virions

Similar to other (+)ssRNA viruses ^31,32^ and depending on the virus production conditions, we observe some level of sgRNA packaging. For low MOI infection and RNase-treated virus stocks, the packaged sgRNA levels were undetectable (Fig. 5D). However, for higher MOI and non-RNase-treated virus stocks, the proportion of sgRNA-containing particles increased to 30–35% for both HAstV1 and MLB2. This could also include sgRNA-mapped defective genomes. We separated concentrated astrovirus particles that contained sgRNAs by sucrose gradient centrifugation to evaluate whether gRNA and sgRNA are packaged into the same or different particles (Fig. 6A). The analysis of individual fractions revealed separation of gRNA- and sgRNA-containing particles, with only gRNA-containing virions being infectious (Fig. 6B-C). This suggests that only one type of RNA (gRNA or sgRNA) can be packaged per virion, consistent with a compact astrovirus structure ^33^ and RNA packaging strategies in other RNA viruses ^34,35^.

**Figure 6.**
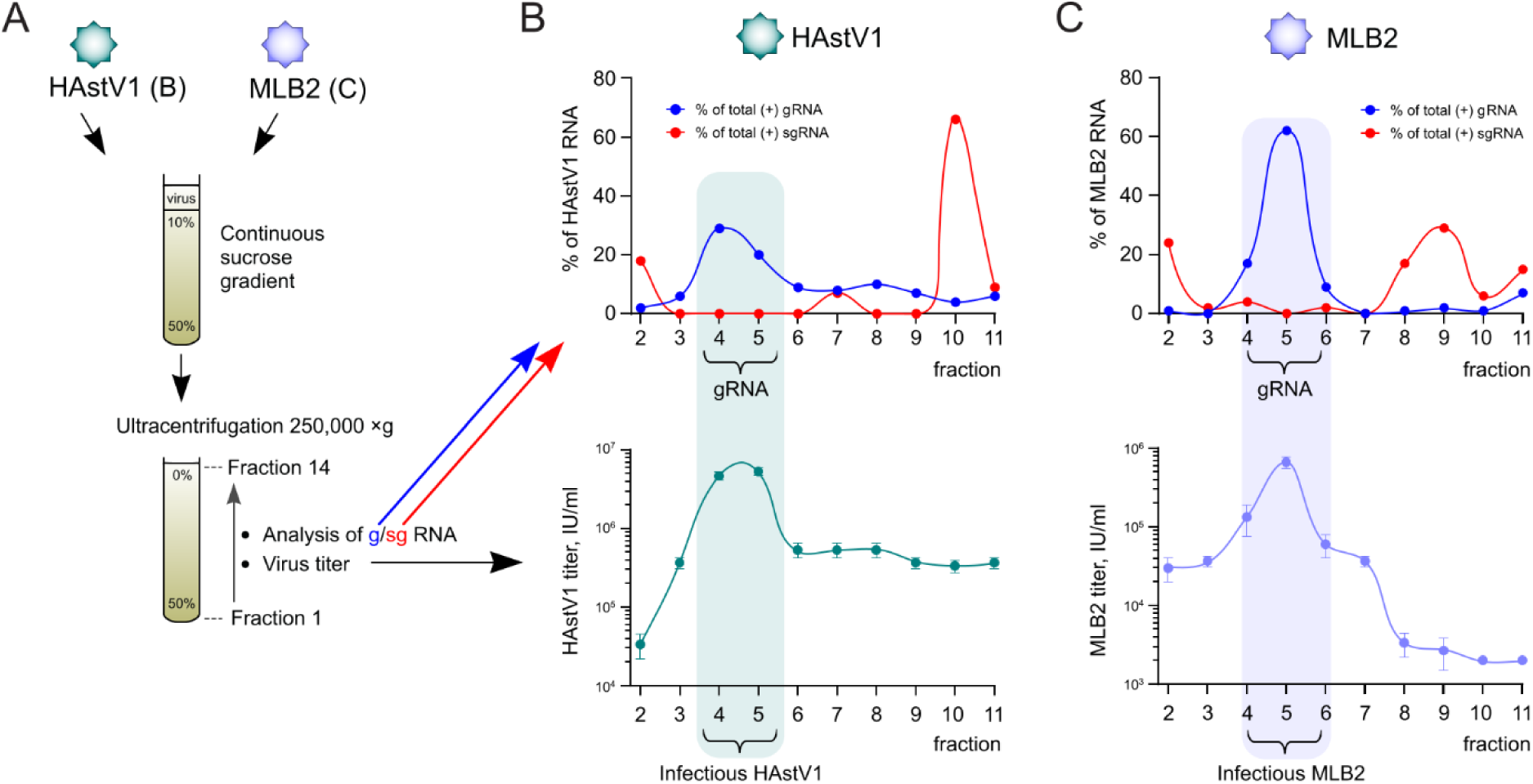
Analysis of (+)gRNAs and (+)sgRNAs in virus particles. (**A**) Experimental setup and analysis of g/sg RNA in fractionated HAstV1 and MLB2 virions. The total amount of both gRNA and sgRNA is set at 100%. (**B, C**) Each fraction was analyzed by ssRT-qPCR (representative top panels) and titrated in Caco-2 (HAstV1) and Huh7.5.1 (MLB2) cells (bottom panels, *n* = 3, mean ± SD). sgRNA levels are calculated from the difference of measured gRNA and g+sgRNA levels.

## Discussion

Replication of the astrovirus RNA genome occurs in intracellular ER membrane-bound vesicles (Fig. 7) ^10,36,37^. Replication complex formation is driven by co-translational membrane targeting through signal peptides located at the N-terminus of the nonstructural polyprotein 1a (nsP1a) ^10,12^. Staining of dsRNA, a hallmark of RNA virus replication, can be detected in perinuclear ER membranes ^10^.

**Figure 7.**
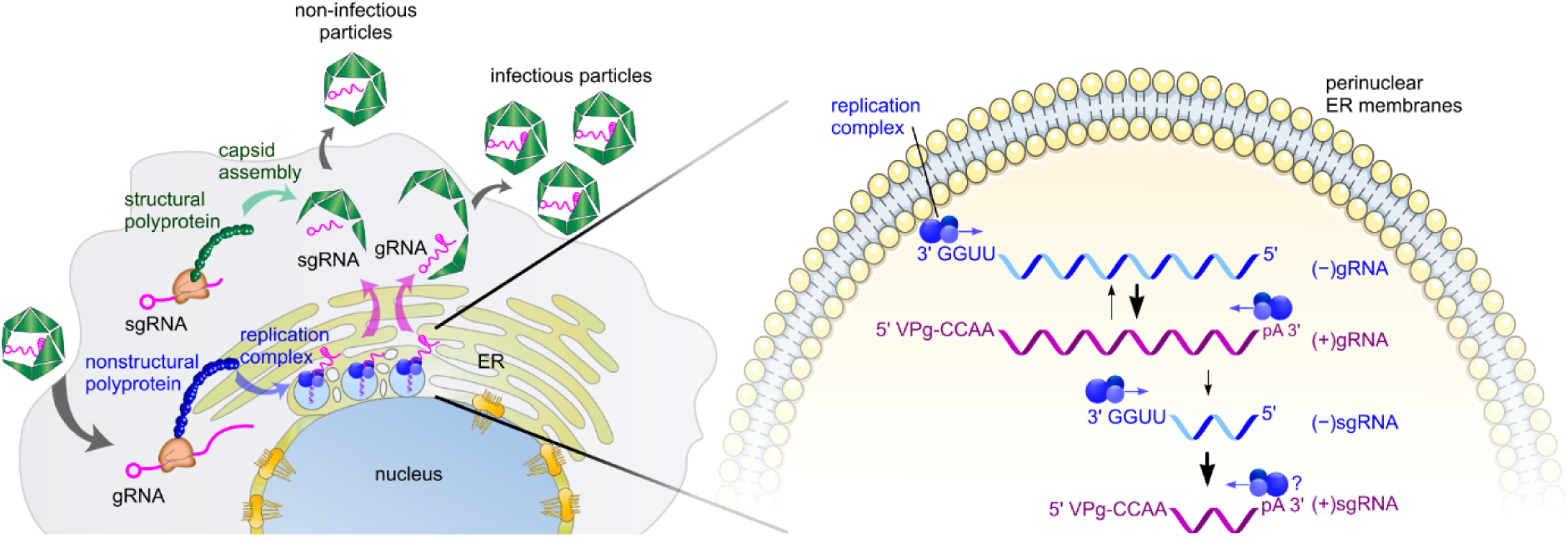
Schematic representation of astrovirus replication. The replication of astroviruses occurs in the perinuclear-derived endoplasmic reticulum (ER) membranes ^10^. RNA replication is represented by four RNA species: (+)gRNA, (+)sgRNA, (−)gRNA, and (−)sgRNA. sgRNA synthesis likely occurs through the premature termination mechanism. The replication balance is shifted towards (+)gRNA and (+)sgRNA synthesis. Packaging preferentially selects (+)gRNA, with packaged (+)sgRNA being non-infectious.

Replication of astrovirus RNA involves the production of both genomic and subgenomic-sized RNA. Synthesis of an sgRNA is conserved in astroviruses, yet the mechanism was previously poorly understood. In contrast, the production of sgRNA is well characterized for many other positive-sense RNA viruses ^18^. Among two sgRNA production mechanisms, internal initiation and premature termination (Fig. 1), the hallmark of the premature termination mechanism is the production of negative-sense sgRNA during virus replication. We demonstrate that astroviruses produce a (−)sgRNA (Fig. 2), thus supporting the premature termination sgRNA production mechanism, rather than the previously suggested internal initiation (Fig. 1) ^23,24^. Formally, it is possible that some or all (−)sgRNAs may be templated from previously produced (+)sgRNAs, either as dead-end accidental products, or for sgRNA amplification. However, the early appearance of (−)sgRNA (Figs 2 and 5), and at quite high levels, suggests that (−)sgRNAs are indeed precursors rather than (only) secondary products.

In our hands, the method of using peaks in histograms of read 5ʹ- or 3ʹ-end mapping positions to identify the termini of transcript species produced variable results – working very well in some cases (e.g. HAstV4) but less well in other cases. We could not assume that the highest peak(s) in a histogram necessarily corresponded to transcript termini. However, with prior expectations of where transcript termini might lie, we could use the technique to provide supporting evidence and to identify termini with single-nucleotide precision in the case of the 3ʹ ends of the negative-sense transcripts where both VPg and a variable-length poly(A) tail are absent. This is further supported by high conservation of gRNA and sgRNA termini (Fig. 3).

The location and function of the subgenomic promoter, alongside functional elements of the premature termination process, are yet to be characterized. Based on our previous studies ^14^, they likely include extended RNA elements located both upstream and downstream of the CCAA sequence in the sgRNA. Likely the upstream elements (acting in a (+)gRNA template) mediate premature termination, whereas the downstream elements (acting at the 3ʹ end of a (−)sgRNA template (or (−)sgRNA:(+)g/sgRNA duplex) mediate RdRp:VPg-primer recruitment. The presence of the conserved 5ʹ-terminal CCAA sequence in gRNAs and sgRNAs implies the same recognition pattern for the RdRp at the 3′ end of the negative strand. Mutating non-VPg-linked CAA nucleotides resulted in reduced gRNA and sgRNA synthesis (Fig. 4). Several other viruses use identical g/sgRNA 5′-terminal recognition motifs, for example, GGUAAU in turnip crinkle virus (*Tombusviridae*) ^38^ and GUGAA in rabbit hemorrhagic disease virus and other caliciviruses (*Caliciviridae*) ^39,40^. The conservation of 5′-terminal gRNA and sgRNA sequences (Fig. 3) suggests that astroviruses use a similar RNA recognition mechanism.

To definitively demonstrate (−)sgRNA as a necessary precursor of (+)sgRNA it would be useful to have a mutant that produces (−)sgRNA but not (+)sgRNA. In other viruses – such as tombusviruses and betanodaviruses – mutation of the first nucleotide of the (+)sgRNA has had this effect ^41,42^. In the case of astroviruses, this mutation (i.e., mutating the first C of the CCAA motif) presumably prevents sgRNA/capsid/virion production, and requires launching from a DNA plasmid as opposed to virus infection as used in Fig. 5. Even with DNase treatment, leftover input DNA prevents useful quantification of viral (−)RNA. Similar studies with other viruses have targeted sgRNAs that are not essential for virion production or have used transcomplementation systems. However, despite substantial attempts, we have not yet succeeded in developing a transcomplementation system for astroviruses.

Taken together, we have identified a (−)sgRNA species ubiquitously present during astrovirus replication, characterized terminal features of astrovirus RNA species, analyzed the dynamics of g/sgRNA production during astrovirus infection, and showed that virions can package sgRNAs but these virions are not infectious. We propose an updated model for astrovirus replication in infected cells, where ER membrane-associated replication ^10^ produces four distinct species of astrovirus RNA, namely (+)/(−)gRNAs and (+)/(−)sgRNAs (Fig. 7). Our findings provide new insights into RNA dynamics during astrovirus infection and open possibilities for astrovirus-specific RNA targeting.

## Materials and Methods

### Cells

BSR cells (single clone of BHK-21 cells) were maintained at 37 °C in DMEM supplemented with 5% fetal bovine serum (FBS), 1 mM L-glutamine and antibiotics. HEK293T cells (ATCC) were maintained in the same media supplemented with 10% FBS. Caco-2 (ATCC) and Huh7.5.1 cells (Apath, Brooklyn, NY) were maintained in the same media supplemented with 10% FBS and non-essential amino acids. All cells were tested mycoplasma negative throughout the work (MycoAlert Mycoplasma Detection Kit, Lonza).

### Virus strains

HAstV1 (pAVIC1, L23513.1), MLB1 (pMLB1, ON398705) and MLB2 (pMLB2, ON398706) were derived from reverse genetics clones ^43,44^. VA1 astrovirus was kindly provided by David Wang (University of St Louis, USA) and clinical HAstV4 was kindly provided by Susana Guix (University of Barcelona, Spain).

### Plasmids and cloning

Dual-luciferase replicons were prepared from the previously described MLB2 ^25^ and HAstV1 ^14,44^ subgenomic replicons. The firefly luciferase gene was inserted in-frame after 16 aa of nsP1a sequence, followed by a 2A separation site and a *SpeI* cleavage site. All mutations were introduced using site-directed mutagenesis and confirmed by sequencing.

### Replicon assays

Linearized replicon-encoding plasmids were used to generate T7 RNAs using HighYield T7 ARCA mRNA Synthesis Kit (Jena Bioscience, RNT-102) according to the manufacturer’s instructions, purified using Zymo RNA Clean & Concentrator kit, and quantified. Cells were transfected in triplicate with Lipofectamine 2000 reagent (Invitrogen), using a previously described reverse transfection protocol ^25^. Three independent experiments, each in triplicate, were performed to confirm the reproducibility of the results.

### Analysis of RNA levels in samples collected from infected cells and virus stocks

Indicated cells grown on 6-well plates (10^6^ cells per well) were infected at MOI 5 or 0.1 and collected at the indicated hpi. RNA from cells (MOI 5) was isolated by Direct-zol RNA MicroPrep (Zymo research); RNA from media-derived samples (MOI 0.1) was isolated using AVL-based (Qiagen, 19073) viral RNA extraction.

The absolute amount of viral RNA was determined by strand-specific RT-qPCR developed based on a previously published technique ^45^. A fixed amount of total RNA was used for reverse transcription (RT) with adaptor-linked strand-specific g/sg primers (Fig. 5A). The qPCR primers were designed to anneal to the adaptor (reverse) and specific (forward) sequences to minimize off-target effects. All primer pairs were validated using T7-transcribed g/sg RNAs. The same transcripts were used for strand-specific quantification.

Reverse transcription was performed using the QuantiTect reverse transcription kit (Qiagen) using the following reverse primers (adaptor is underlined):

HAstV1-SG-RTpos: <u>ACGGATGCGATGTAGAGTGCC</u>CAGTACCATTAACAGCAGATGC
HAstV1-G-RTpos: <u>ACGGATGCGATGTAGAGTGCC</u>GCACGGGCTCAAGGTTTTCGTTAAC
HAstV1-SG-RTneg: <u>GGCCGTCATGGTGGCGAATAA</u>GGAAGCACTCAGTTTGGCCCTGTG
HAstV1-G-RTneg: <u>GGCCGTCATGGTGGCGAATAA</u>CTATGTATAGAGACATCTTTGGCATG
MLB2-SG-RTpos: <u>ACGGATGCGATGTAGAGTGCC</u>GAATTCAACGTAGGGCCAGTTG
MLB2-G-RTpos: <u>ACGGATGCGATGTAGAGTGCC</u>GGTGCAGGTCCTTTCTTAGTATTTG
MLB2-SG-RTneg: <u>GGCCGTCATGGTGGCGAATAA</u>CATTATGGAGACTTAAGTGGCTGGAG
MLB2-G-RTneg: <u>GGCCGTCATGGTGGCGAATAA</u>TAGCCATGTATAAAGATGTCTTTGG

Quantitative PCR was performed in triplicate using SsoFast EvaGreen Supermix (Bio-Rad) in a ViiA 7 Real-time PCR system (Applied Biosystems) for 40 cycles with two steps per cycle using the following qPCR primers:

SS-pos-Rq: ACGGATGCGATGTAGAGTGCC
HAstV1-pos-SG-Fq: GGAAGCACTCAGTTTGGCCCTG
HAstV1-pos-G-Fq: CTATGTATAGAGACATCTTTGGCATGTG
SS-neg-Rq: GGCCGTCATGGTGGCGAATAA
HAstV1-neg-SG-Fq: CAGTACCATTAACAGCAGATGC
HAstV1-neg-G-Fq: GCACGGGCTCAAGGTTTTCGTTAAC
MLB2-pos-SG-Fq: CATTATGGAGACTTAAGTGGCTGGAGG
MLB2-pos-G-Fq: TAGCCATGTATAAAGATGTCTTTGGAATG
MLB2-neg-SG-Fq: GAATTCAACGTAGGGCCAGTTG
MLB2-neg-G-Fq: GGTGCAGGTCCTTTCTTAGTATTTG

### Analysis of RNA by RNA-seq

Caco-2 cells were infected at MOI 5 with HAstV1, HAstV4 or VA1 and incubated for 24 h (additionally 6, 12 and 18 h for HAstV1). Huh7.5.1 cells were infected at MOI 5 with MLB1 or MLB2 and incubated for 24 h. The total RNA was extracted by Direct-zol RNA Prep kit (Zymo research). RNA was fragmented, and 50–150 nt fragments were used for RNA-seq library preparation using an Illumina library protocol ^46,47^, except for Ribo-Zero rRNA removal kit (Illumina) was used to deplete ribosomal RNA. Amplicon libraries were deep sequenced using an Illumina NextSeq platform. For the indicated experiments, proteinase K (PK, NEB) treatment was performed using 10 µg of total RNA extracted from infected cells, 1% SDS (final concentration) at 42°C for 45 minutes. We avoided poly(A) selection for the RNA-seq as this would bias RNA-seq density towards the 3′ ends of transcripts.

### Analysis of high throughput sequencing datasets

Libraries for the first high throughput sequencing experiment (HAstV1; 6, 12, 18 hpi timecourse) were prepared without “Unique Molecular Identifiers” (UMIs), and sequenced on an Illumina NextSeq platform using single-end sequencing with 76 cycles. Read quality was assessed with the FASTX-Toolkit (http://hannonlab.cshl.edu/fastx_toolkit) but, based on the results, no quality filtering was applied. Adaptor sequences were trimmed using fastx_clipper (FASTX-Toolkit) with parameters -Q33 -a TGGAATTCTCGGGTGCCAAGGAACTCCAGTCA -l 30 -c -n -v. The “-c” option discards reads where the adaptor sequence was not found and the -l 30 option discards trimmed reads <30 nt in length. For the single-end sequencing (in contrast to the paired-end sequencing used later), use of the -c option is important because we are interested in the exact location of the 3′ end of reads (especially for the (−)RNA reads).

Trimmed reads were mapped to *Homo sapiens* rRNA (NCBI accessions NR_003287.2, NR_023379.1, NR_003285.2 and NR_003286.2), the HAstV1 genome (NCBI accession L23513.1, with point changes U2191C, A2335G, A2716U, U2719C, G2936A, A3213G and U6010C, and edited to have exactly 50 nt of poly(A) tail), and a *H. sapiens* mRNA database (35,809 RefSeq mRNAs; downloaded from NCBI 24 Jan 2013) using bowtie1 ^48^ in single-end mode, with parameters -v 2 --best (i.e. maximum two mismatches; report best match). The full set of reads was mapped to each database. The 50-nt poly(A) tail was added to the HAstV1 genome to allow (most) *bona fide* virus reads overlapping into the poly(A) tail to be successfully mapped. However, it also allows A*n* sequences derived from host mRNA poly(A) tails to mismap to the virus bowtie database. Therefore, we subsequently removed potentially non-specific poly(A)-mapped reads by discarding vRNA-mapped reads that contained fewer than 20 nt of vRNA sequence upstream of the poly(A) tail; specifically, we discarded (+)vRNA-mapped reads whose 5′ end mapped <20 nt upstream of the poly(A) start site and, for symmetry, we also discarded (−)vRNA-mapped reads where the 5′ end of the reverse complement of the read mapped <20 nt upstream of the poly(A) start site.

Libraries for the second high throughput sequencing experiment (HAstV1; 18 and 24 hpi; with or without proteinase K treatment; etc) were prepared with 7-nt random sequence tags (UMIs) at the 5′ of each read, and sequenced on an Illumina NextSeq platform using paired-end sequencing with 76 cycles in each direction, and obtained from the provider with 3′ adaptor sequences already trimmed. In this set of experiments, we explored using two different gel slices: nominally 75–150 nt and 120–150 nt reads, but in fact this distinction was not apparent in the observed length distribution of virus-mapped reads (Fig. S21); therefore, the two gel slices were considered as technical repeats. Read quality was assessed with the FASTX-Toolkit and was quite variable across the libraries. To achieve higher quality uniformly across all libraries, we trimmed all reads to 30 nt (R2; reverse reads) and 37 nt (R1; forward reads) using fastx_trimmer (FASTX-Toolkit), where the 5′-terminal 7 nt of the R1 reads are the 7-nt UMIs, removed before mapping. Read pairs where either R1 or R2 had already been adaptor trimmed to <37 nt or <30 nt, respectively, were discarded. Next, we discarded read pairs where the 37-nt trimmed R1 read contained any ‘N’s, deduplicated by saving only one occurrence of any group of read pairs with an identical R1 sequence (including the 5′-terminal 7-nt UMI), and removed the UMI from R1. The remaining 30-nt R1 and R2 reads were mapped to the above-described *H. sapiens* rRNA, HAstV1, and *H. sapiens* mRNA databases using bowtie1 in paired-end mode, with parameter -v 2. Bowtie1 only reports matches where both R1 and R2 have syntenous alignments, and the corresponding inferred fragment size is from 30 to 250 nt. Potentially non-specific virus poly(A)-mapped read pairs were removed as described above. Across the 16 libraries, the mean and standard deviation of the inferred fragment size for virus mapped read pairs ranged 58–75 nt and 8.5–14.1 nt, respectively. Unusually large inferred fragment sizes (e.g. >115 nt) likely derive from defective transcripts with internal deletions. Although a very small fraction of the total counts, these inferred fragments may contaminate the virus genome coverage plots. Therefore, for each of the 16 libraries, we removed read pairs where the inferred fragment length was >3 standard deviations above the mean for the respective library.

Libraries for the third high throughput sequencing experiment (HAstV4, MLB1, MLB2, VA1) were prepared without UMIs, and sequenced on an Illumina NextSeq platform using paired-end sequencing with 37 cycles in the forward direction and 38 cycles in the reverse direction. Read quality was assessed with the FASTX-Toolkit and, as above, R1 and R2 reads were trimmed to 30 nt. In the absence of UMIs, deduplication was not performed. In contrast to the second experiment, read pairs with ‘N’s in R1 were not removed (as this step was only implemented above to make deduplication more robust), but in any case only 0.060% of 30-nt trimmed R1 reads and 0.031% of 30-nt trimmed R2 reads contained ‘N’s. The 30-nt R1 and R2 reads were mapped to the *H. sapiens* rRNA and *H. sapiens* mRNA databases, besides the relevant viral genome (HAstV4, MLB1, MLB2 or VA1) using bowtie1 in paired-end mode, with parameter -v 2. NCBI accessions were as follows: HAstV4 – PX619609; MLB1 – ON398705.1 with a 30-nt deletion at 5908–5937 as previously described ^25^; MLB2 – ON398706.1 with a U3112C point change and a 5-nt deletion at 5840–5844 as previously described ^25^; and VA1 – FJ973620.1 with point changes C499U, A2672G, U5162C, C5369U, A5470U; in each case sequences were modifed to have exactly 50 nt of poly(A) tail. As described above, potentially non-specific virus poly(A)-mapped reads were removed, and read pairs where the inferred fragment length was >3 standard deviations above the mean for the respective library were also removed.

In the following, mapped ‘fragments’ refers to mapped full-length single-end sequencing reads or inferred fragments based on mapped 30-nt R1/R2 read pairs. Normalization for library size was based on the number of fragments mapped to *H. sapiens* (+)mRNA plus (+)vRNA. Relative densities for gRNA-only and gRNA/sgRNA-overlap regions were calculated from the number of fragments whose 5′ end mapped from nucleotide 1 to 1 nt upstream of the sgRNA start site, or from the sgRNA start site to 115 nt less than the genome length excluding the poly(A) tail, respectively. For fragments derived from (−)RNA, we used the 5′ end of the reverse complement of the fragment. The poly(A) tail was excluded because it is of unknown and variable length. The 115 nt 3′ buffer was used because, across all libraries, after the above processing, the maximum length of a virus mapped fragment was 115 nt. The sgRNA start site was based on a 5′-CCAA location corresponding to that annotated in Fig. 3C (viz. nt 4315, 4313, 3830, 3830 and 4199 for HAstV1, HAstV4, MLB1, MLB2 and VA1, respectively). For the bargraphs, counts were normalized by the length (in kb) of the respective gRNA-only or gRNA/sgRNA-overlap region to obtain ‘reads per kb per million mapped reads’ values. For the ‘relative density’ bar graphs, these values are simply scaled so that the sum of (+)gRNA, (+)sgRNA, (−)gRNA and (−)sgRNA is unity for each library.

Virus-derived reads containing untemplated nucleotides may fail to map with bowtie1. Therefore, to test for the possibility of untemplated nucleotides added at the start of the (−)gRNA and/or (−)sgRNA, we looked at the 5′ ends of (+)RNA-sense R2 reads (such reads derive from (−)RNA and their 5′ ends correspond to the 3′ ends of (−)RNA fragments). We searched the processed (as described above) but unmapped R2 read files using nucleotides 6–20 of the (+)gRNA and (+)sgRNA as 15-nt queries, extracted all matches, truncated matches at the start of the query, and sorted these truncated matches by abundance for each library. For the HAstV1 paired-end sequencing, R2 matches to the gRNA query (AGGGGGGUGGUGAUU) were found for 13 of the 16 libraries, and in all 13 libraries the most abundant upstream nucleotides were the expected CCAAG (141 occurrences in total) corresponding to the gRNA 5′ terminus. Excluding singletons, the only other upstream sequences were truncations of CCAAG (i.e. CAAG, AAG, AG and G). For R2 matches to the sgRNA query (GAAGUGUGAUGGCUA), the most abundant upstream sequence in each library was usually the expected CCAAG (12 of 16 libraries; mean 21 occurrences per library). The four exceptions were all low abundance in the respective library (1 × GGGAGGACCAAA; 2 × AGGGGAGGACCAAA; 1 × GGGAGGACCAAA; 1 × GGGGAGGACCAAA) and all map to gRNA. Excluding singletons, the only other represented sequences were the gRNA- mapping GAGGGGAGGACCAAA and all possible 5′ truncations thereof down to A. Thus we found no evidence for untemplated nucleotides in HAstV1. We performed similar analyses for the HAstV4, MLB1, MLB2 and VA1 paired-end datasets and again found no evidence for untemplated nucleotides at the 3′ end of the (−)gRNA or (−)sgRNA. For gRNA, the read counts were too low for the MLB1 and MLB2 datasets, with only one gRNA 5′-proximal-query-containing R2 read for each (albeit, both with the expected 5′-terminal CCAAG). For sgRNA, exceptions to the expected CCAAA as the most abundant 5′-terminal sequence in 5′-proximal-query-containing R2 reads were MLB2 rep. 1 (no reads found), and MLB1 rep. 2 (2 × GGGUGGACCAAA) and VA1 rep. 2 (1 × GGUCCAAA), both of which map to gRNA. Ignoring singletons, none of the extracted sequences had non-templated nucleotides.

As noted in the main text, for HAstV1, HAstV4 and VA1, the *bona fide* 3′ end of the (−)gRNA could be determined from a peak in the histogram of RNA-seq 3′-end mapping positions, and in all cases is 3′-GGUU, corresponding to an inferred 5′-CCAA terminus for the (+)gRNA. However, for MLB1 and MLB2, there were too few (−)RNA reads for this analysis. We performed *de novo* assemblies of the MLB1 and MLB2 libraries using Trinity v. 2.2.0 ^49^. Our two MLB1 assemblies had 5′-CCAAG and 5′-gcgccCCAAG, respectively, with the latter assumed to involve a spurious addition of 5 nt. For our two MLB2 libraries, the assemblies both had 5′-terminal CAAG (i.e. lacking one of the two expected 5′-terminal ‘C’s). Our MLB1 and MLB2 assemblies have 99.95% and 99.84% nucleotide identity to NCBI accessions MK089434 and MK089435, respectively, both of which have 5′-terminal CCAAG. Therefore we infer that the correct 5′-terminus of both MLB1 and MLB2 (+)gRNAs is most likely to be CCAAG.

The sequence logo (Fig. 3B) was produced via the Weblogo server at https://weblogo.berkeley.edu/logo.cgi.

### Statistical analyses

Data were graphed and analyzed using GraphPad Prism. Significance values are shown as \*\*\*\**p* < 0.0001, \*\*\**p* < 0.001, \*\**p* < 0.01, \**p* < 0.05, ns – non-significant.

## Data availability

The sequencing data reported in this paper have been deposited in ArrayExpress (http://www.ebi.ac.uk/arrayexpress) under the accession number E-MTAB-16460.

## Acknowledgments

This work was funded by a Sir Henry Dale Fellowship [220620/Z/20/Z] from the Wellcome Trust and the Royal Society to VL. A.E.F. is supported by a Wellcome Trust Senior Research Fellowship [220814/Z/20/Z]. DN is supported by the Elizabeth Mann-funded studentship from the Department of Pathology. The authors thank members of the Lulla lab for helpful discussions. We thank Cambridge Genomic Services for high throughput sequencing.

For the purpose of Open Access, the authors have applied a CC BY public copyright license to any Author Accepted Manuscript (AAM) version arising from this submission.

The authors declare that they have no conflict of interest.

## Notes

### Competing Interest Statement

The authors have declared no competing interest.

