## Supplementary information for "The dynamics and strategy of RNA replication in astroviruses"

**Supplementary Figure 1. Total coverage of vRNA(+) and vRNA(−).** Caco-2 cells were infected with HAstV1 at MOI 5 and harvested at 6, 12 or 18 hpi in duplicate. Fragments were mapped to vRNA(+) or vRNA(−), and total coverage summed. The y-axis scale is arbitrary but vRNA(−) coverage is scaled relative to vRNA(+) coverage by the indicated factor to aid visualization.

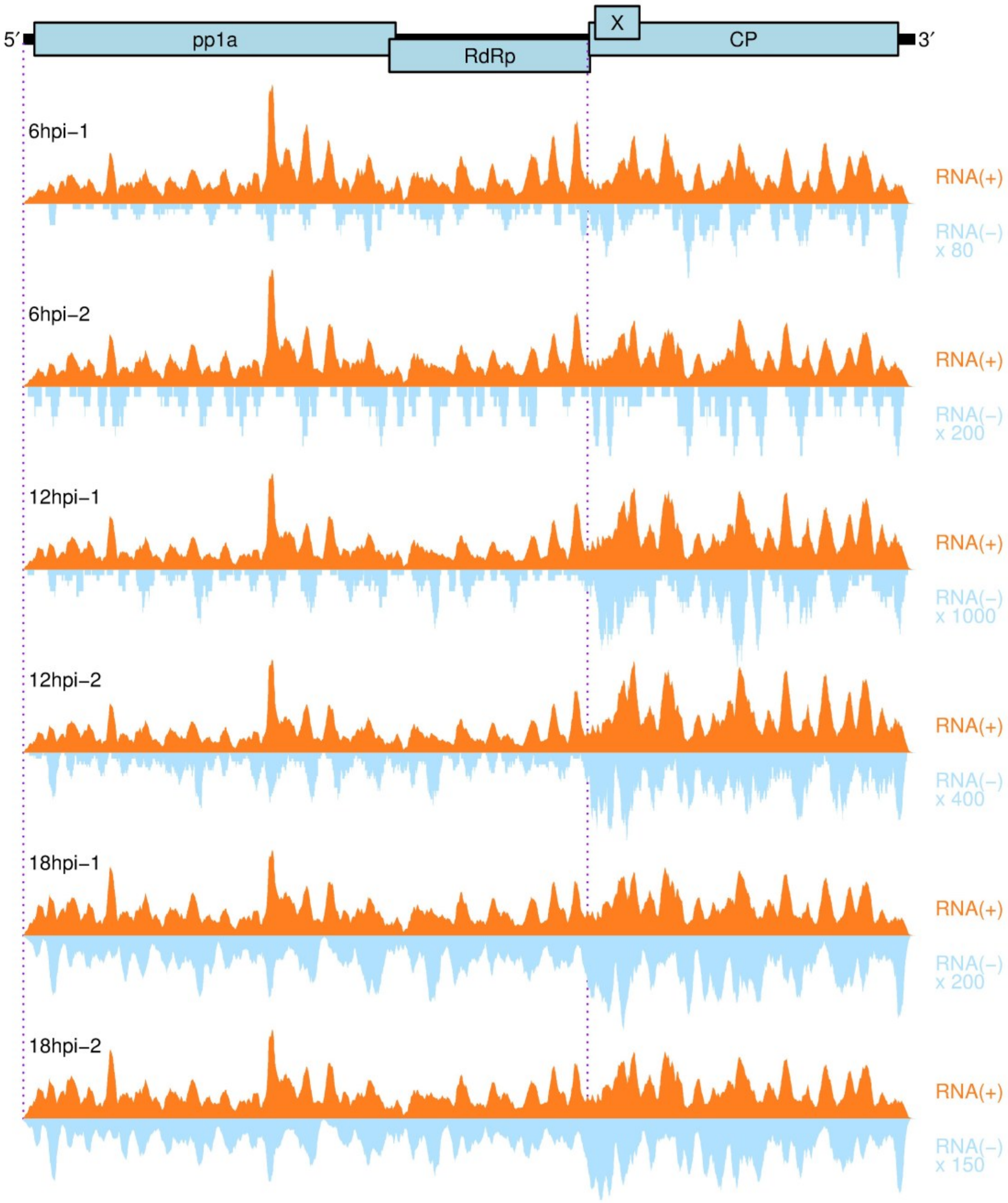

**Supplementary Figure 2. Histograms showing positions of 5′ ends of fragments mapping to vRNA(+).** Caco-2 cells were infected with HAstV1 at MOI 5 and harvested at 6, 12 or 18 hpi in duplicate. Counts are normalized to fragments per million fragments mapped to vRNA(+) or host mRNA(+) (RPM).

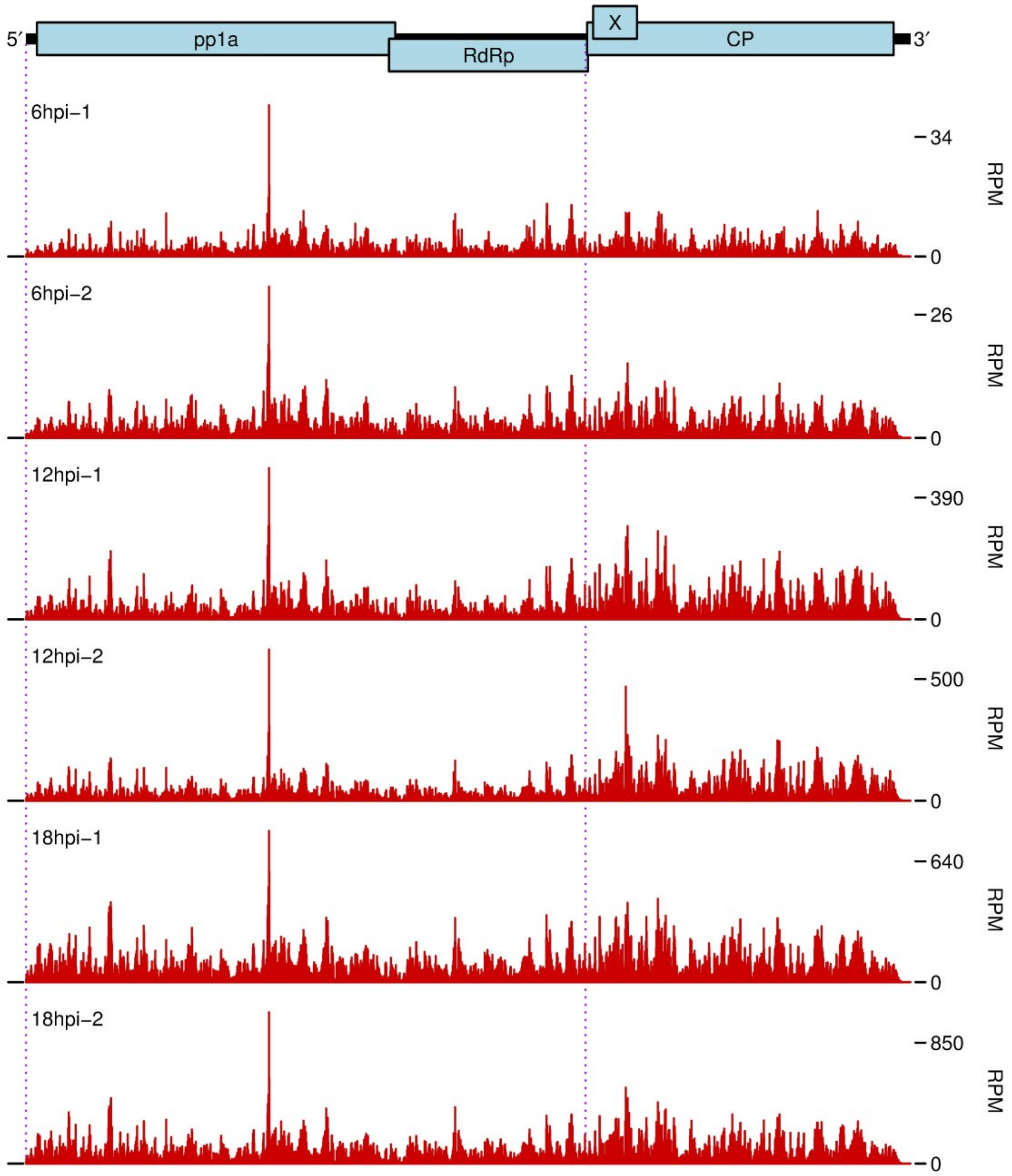

**Supplementary Figure 3. Histograms showing positions of 3′ ends of fragments mapping to vRNA(−).** Caco-2 cells were infected with HAstV1 at MOI 5 and harvested at 6, 12 or 18 hpi in duplicate. Counts are normalized to fragments per million fragments mapped to vRNA(+) or host mRNA(+) (RPM). Histograms show 3′ ends of negative-sense fragments, corresponding to 5′ ends of the positive-sense reverse complements of the fragments.

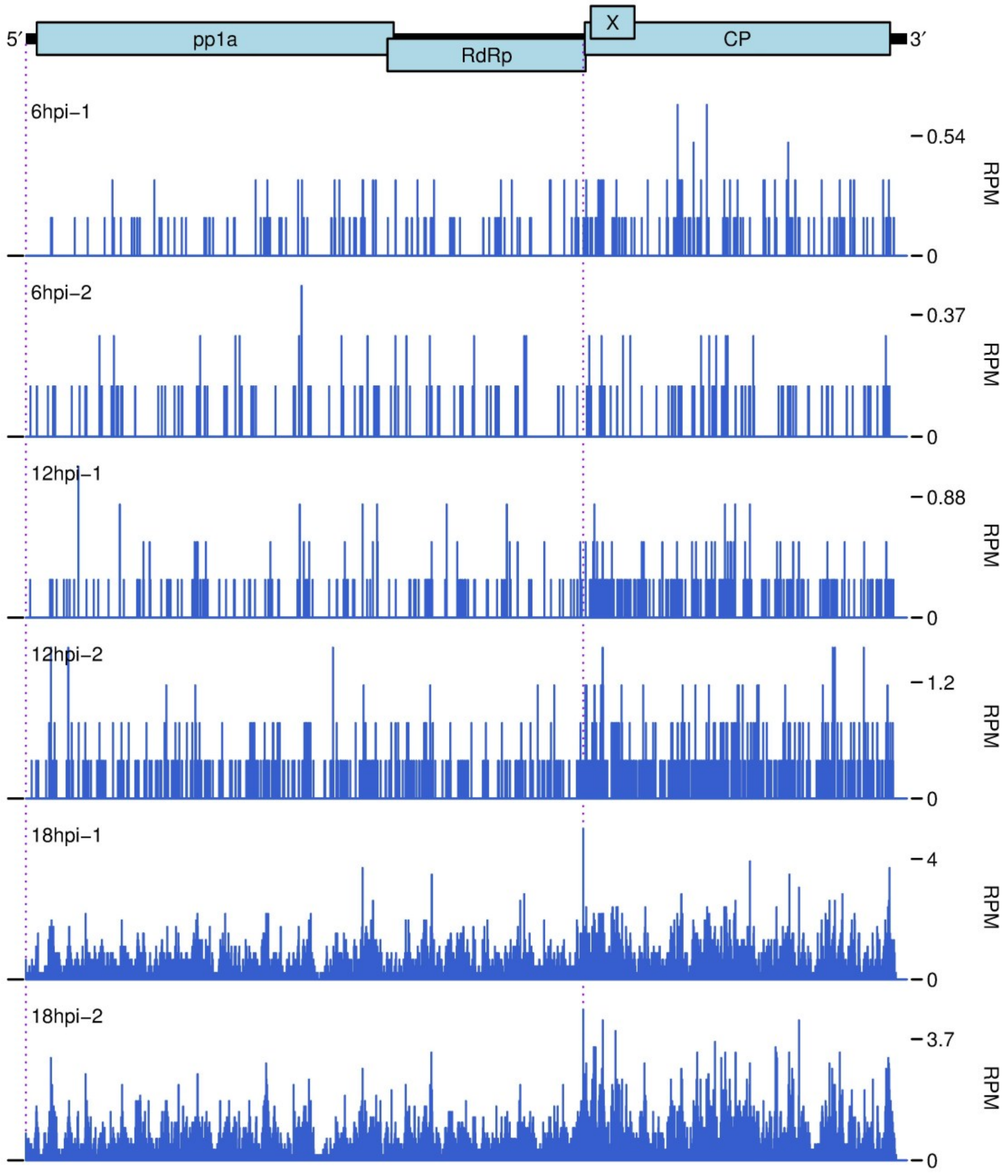

**Supplementary Figure 4. Histograms showing positions of 5′ ends of fragments mapping to vRNA(+).** Caco-2 cells were infected with HAstV1 at MOI 5 and harvested at 18 or 24 hpi in duplicate, with or without proteinase K (PK) treatment; nominally 75–150 nt fragments were selected for sequencing. Counts are normalized to fragments per million fragments mapped to vRNA(+) or host mRNA(+) (RPM).

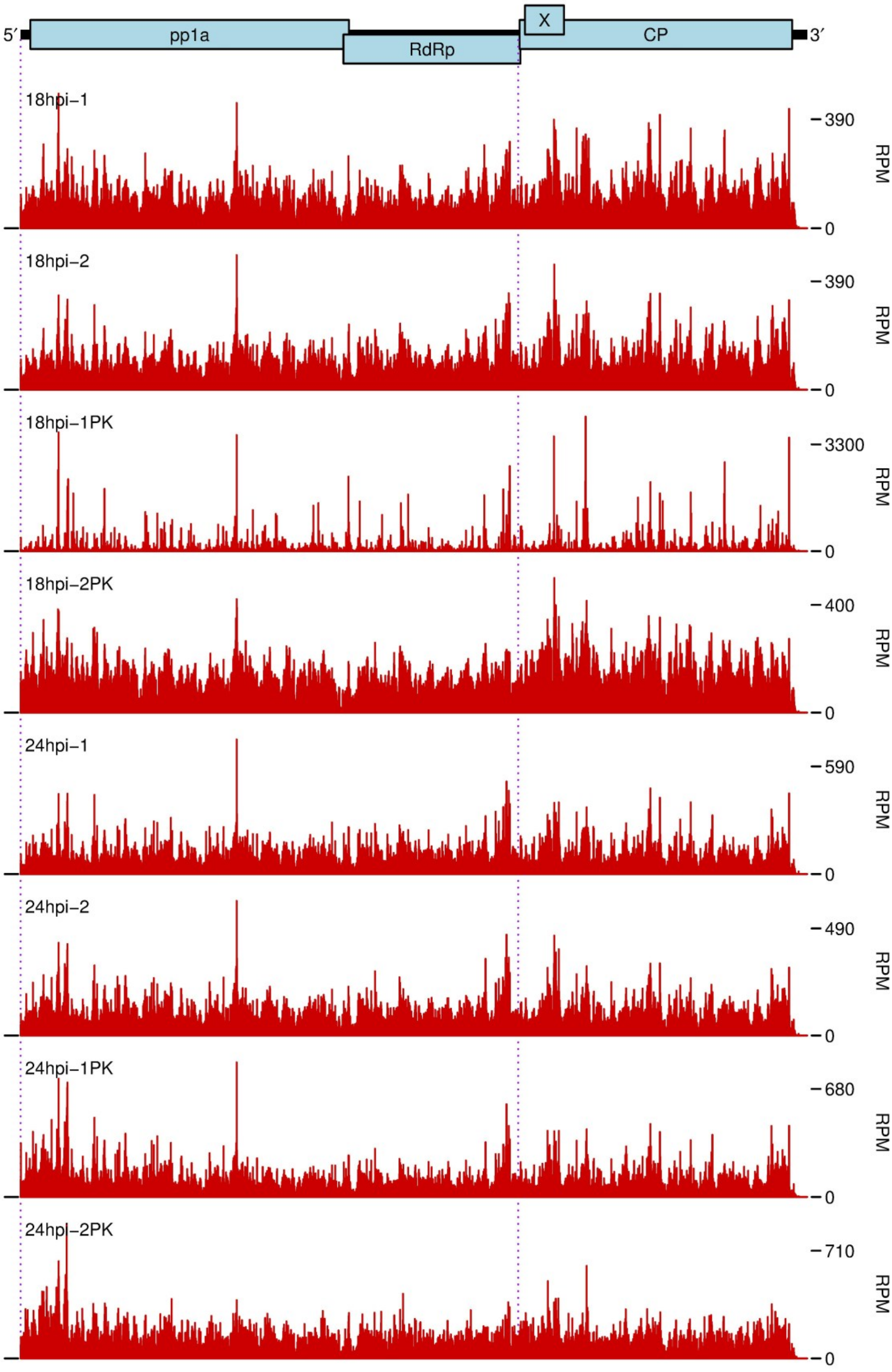

**Supplementary Figure 5. Histograms showing positions of 5′ ends of fragments mapping to vRNA(+).** Caco-2 cells were infected with HAstV1 at MOI 5 and harvested at 18 or 24 hpi in duplicate, with or without proteinase K (PK) treatment; nominally 120–150 nt fragments were selected for sequencing. Counts are normalized to fragments per million fragments mapped to vRNA(+) or host mRNA(+) (RPM).

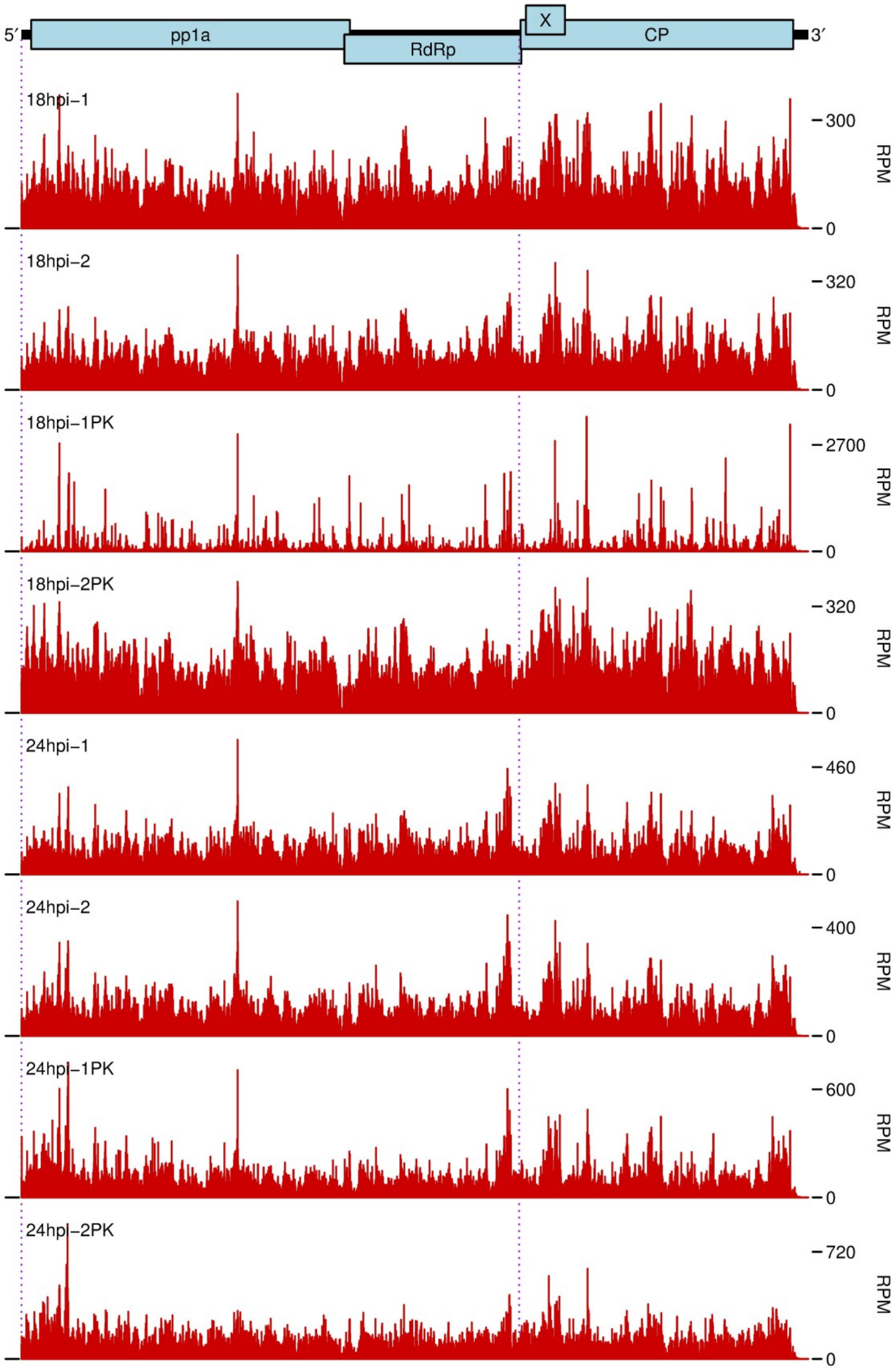

**Supplementary Figure 6. Levels of astrovirus positive/negative-sense gRNA/sgRNA species.** Caco-2 cells were infected with HAstV1 at MOI 5 and harvested at 18 or 24 hpi in duplicate, with or without proteinase K (PK) treatment. (**A**) Bar graphs showing the density of mapped fragments in the sgRNA region (pink), outside of the sgRNA region (red) and the difference (yellow). Coverage was quantified as fragments per kilobase per million fragments mapped to vRNA(+) or host mRNA(+). Fragments mapping to the sgRNA region may derive from either gRNA or sgRNA; the difference (yellow) in density between the sgRNA and non-sgRNA regions was used to estimate the relative abundance of sgRNA, whereas the density in the non-sgRNA region (red) was used to estimate the relative abundance of gRNA. (**B**) Relative densities of (+)gRNA, (+)sgRNA, (−)gRNA and (−)sgRNA. Numbers below bars show the estimated sgRNA:gRNA ratio (1 d.p.). (**C**) Estimated (−):(+) ratio for gRNA and sgRNA species. The (−):(+) sgRNA ratios are not shown for the 24 hpi samples because the (+)sgRNAvalues are unreliable at this time point (insufficient RNA-seq density difference between the gRNA/sgRNA overlap region and the gRNA-only region).

**A**

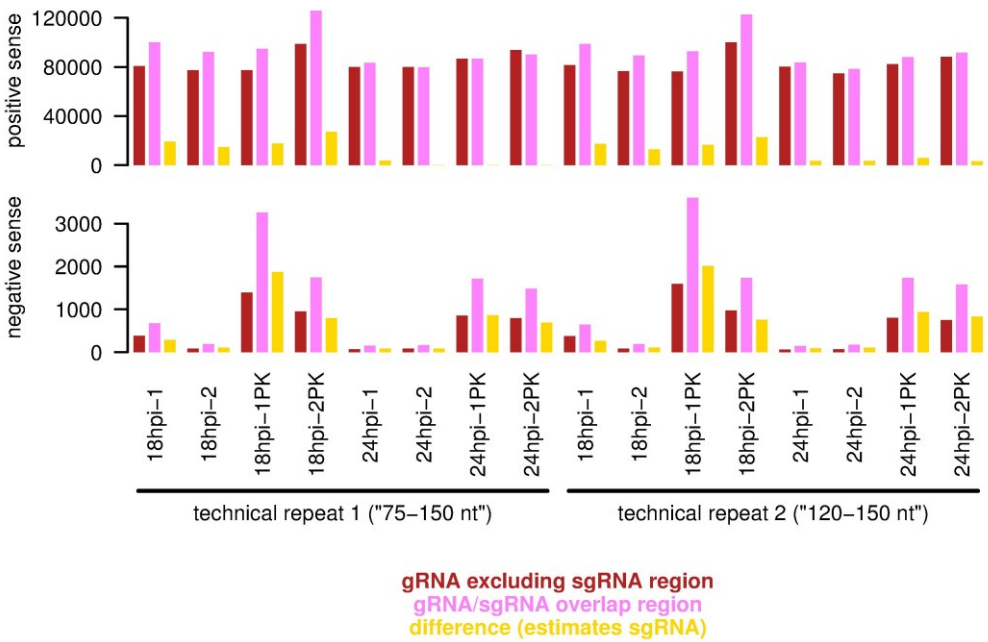


**B**

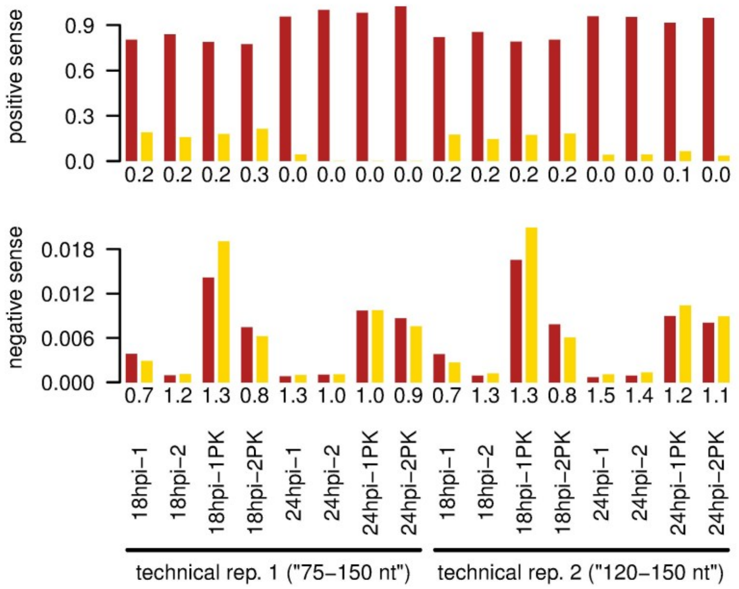


**C**

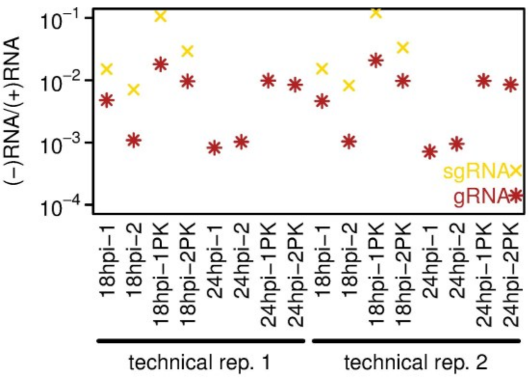

**Supplementary Figure 7. Total coverage of vRNA(+) and vRNA(−).** Caco-2 cells were infected with HAstV1 at MOI 5 and harvested at 18 or 24 hpi in duplicate, with or without proteinase K (PK) treatment; nominally 75–150 nt fragments were selected for sequencing. Fragments were mapped to vRNA(+) or vRNA(−), and total coverage summed. The y-axis scale is arbitrary but vRNA(−) coverage is scaled relative to vRNA(+) coverage by the indicated factor to aid visualization.

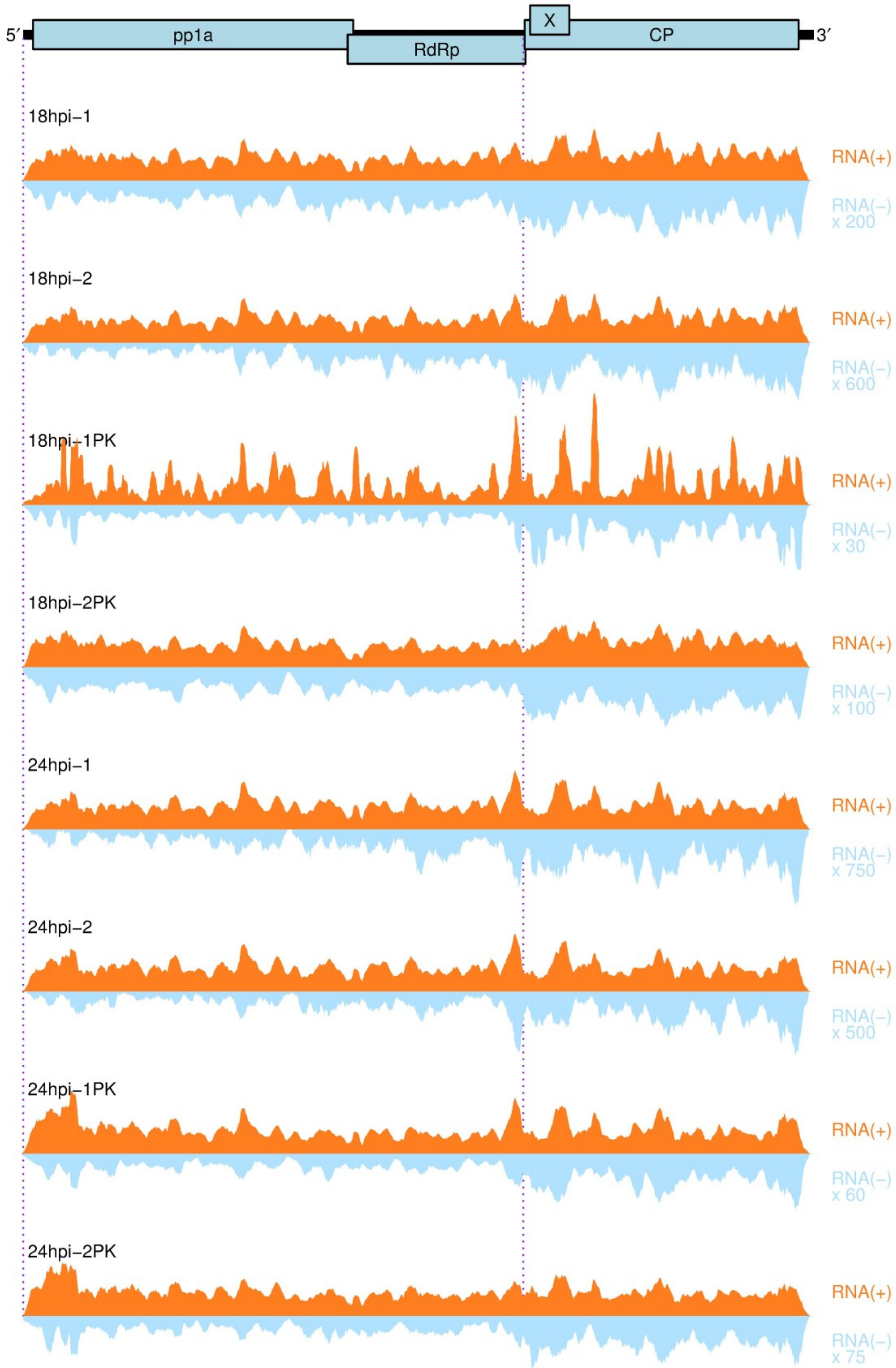

**Supplementary Figure 8. Total coverage of vRNA(+) and vRNA(−).** Caco-2 cells were infected with HAstV1 at MOI 5 and harvested at 18 or 24 hpi in duplicate, with or without proteinase K (PK) treatment; nominally 120–150 nt fragments were selected for sequencing. Fragments were mapped to vRNA(+) or vRNA(−), and total coverage summed. The y-axis scale is arbitrary but vRNA(−) coverage is scaled relative to vRNA(+) coverage by the indicated factor to aid visualization.

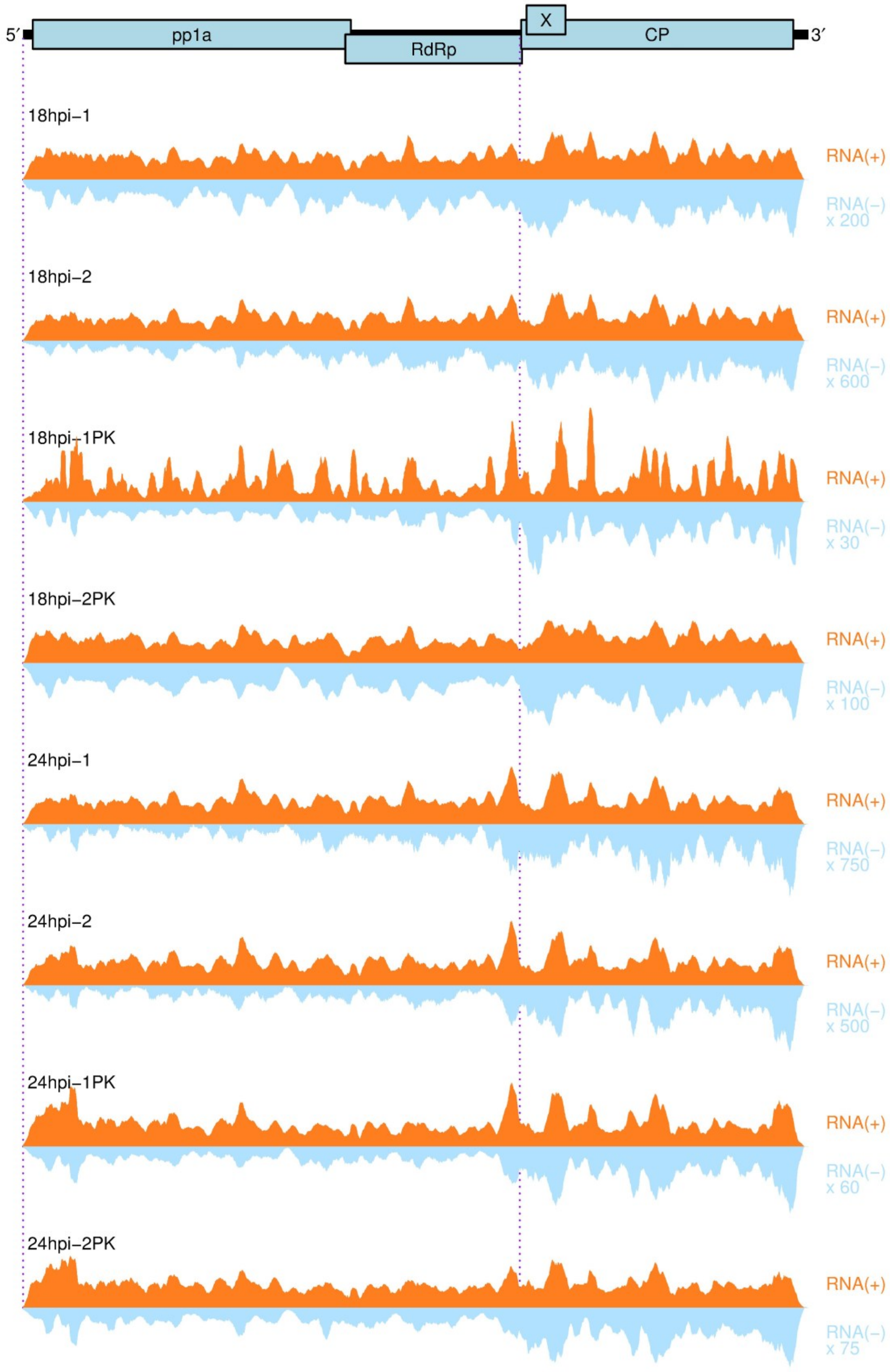

**Supplementary Figure 9. Histograms showing positions of 3′ ends of fragments mapping to vRNA(−).** Caco-2 cells were infected with HAstV1 at MOI 5 and harvested at 18 or 24 hpi in duplicate, with or without proteinase K (PK) treatment; nominally 75–150 nt fragments were selected for sequencing. Counts are normalized to fragments per million fragments mapped to vRNA(+) or host mRNA(+) (RPM). Histograms show 3′ ends of negative-sense fragments, corresponding to 5′ ends of the positive-sense reverse complements of the fragments.

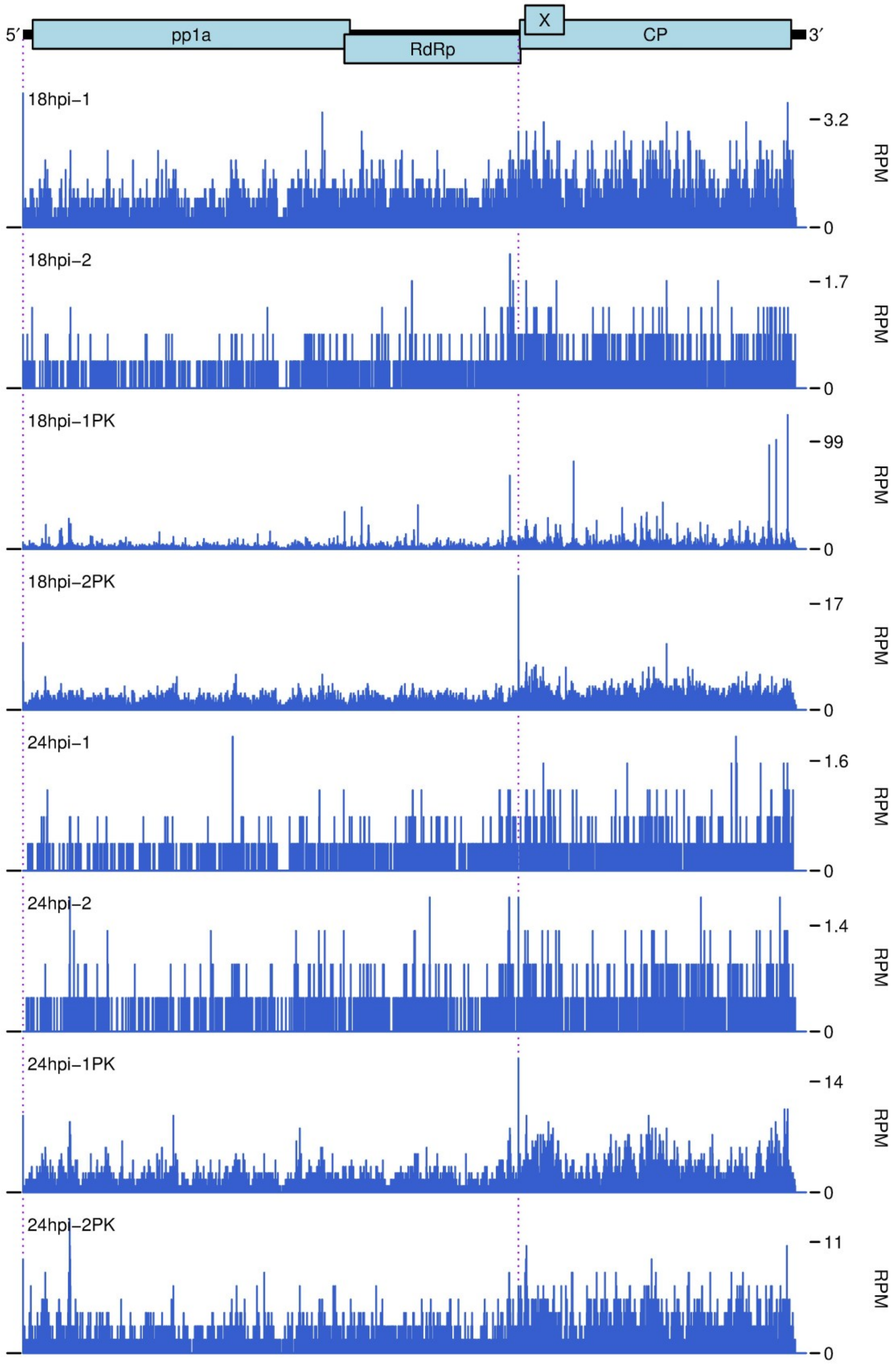

**Supplementary Figure 10. Histograms showing positions of 3′ ends of fragments mapping to vRNA(−).** Caco-2 cells were infected with HAstV1 at MOI 5 and harvested at 18 or 24 hpi in duplicate, with or without proteinase K (PK) treatment; nominally 120–150 nt fragments were selected for sequencing. Counts are normalized to fragments per million fragments mapped to vRNA(+) or host mRNA(+) (RPM). Histograms show 3′ ends of negative-sense fragments, corresponding to 5′ ends of the positive-sense reverse complements of the fragments.

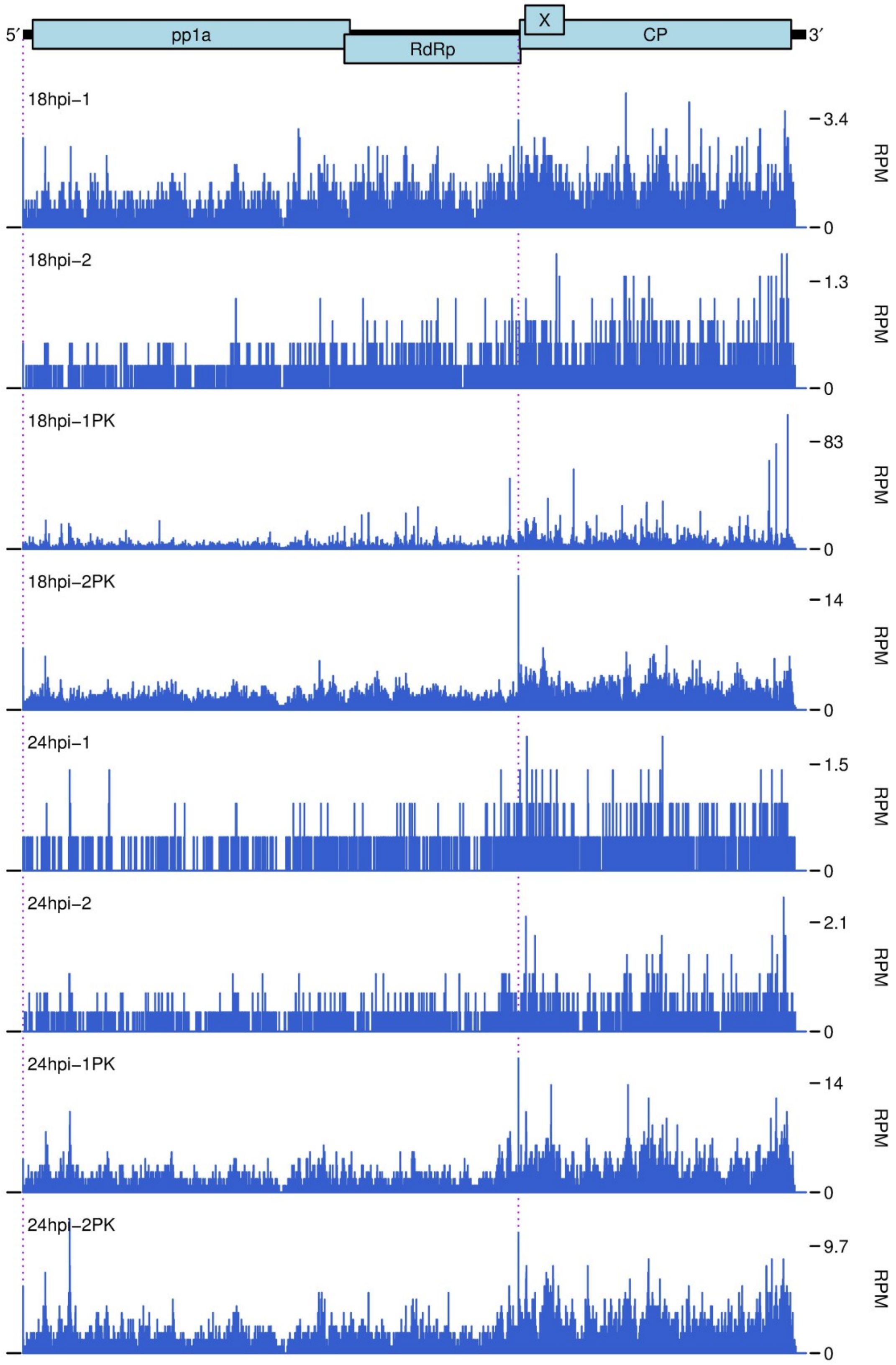

**Supplementary Figure 11. Levels of astrovirus positive/negative-sense gRNA/sgRNA species.** Cells were infected with HAstV4 (in quadruplicate), or MLB1, MLB2 or VA1 (in duplicate) viruses at MOI 5 and harvested at 24 hpi Caco-2 cells were used for HAstV4 and VA1, and Huh7.5.1 cells were used for MLB1 and MLB2. **(A)** Bar graphs showing the density of mapped fragments in the sgRNA region (pink), outside of the sgRNA region (red) and the difference (yellow). Coverage was quantified as fragments per kilobase per million fragments mapped to vRNA(+) or host mRNA(+). Fragments mapping to the sgRNA region may derive from either gRNA or sgRNA; the difference (yellow) in density between the sgRNA and non-sgRNA regions was used to estimate the relative abundance of sgRNA, whereas the density in the non-sgRNA region (red) was used to estimate the relative abundance of gRNA. (**B**) Relative densities of (+)gRNA, (+)sgRNA, (−)gRNA and (−)sgRNA. Numbers below bars show the estimated sgRNA:gRNA ratio (1 d.p.). (**C**) Estimated (−):(+) ratio for gRNA and sgRNA species.

**A**

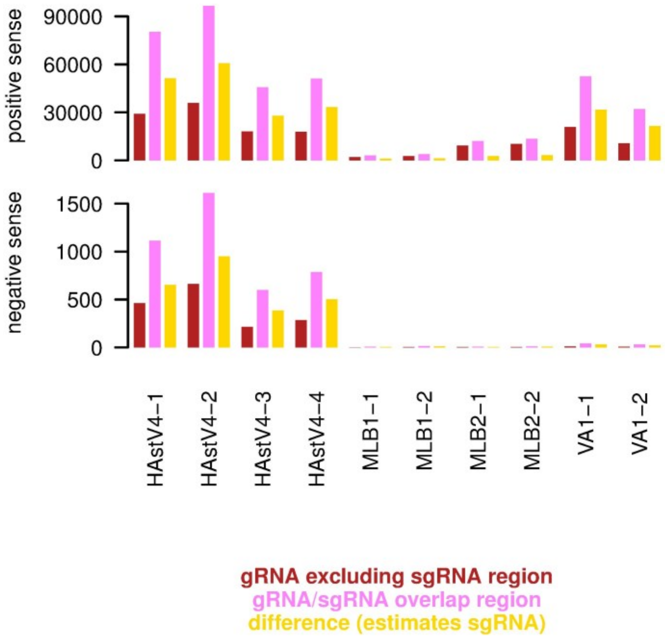


**B**

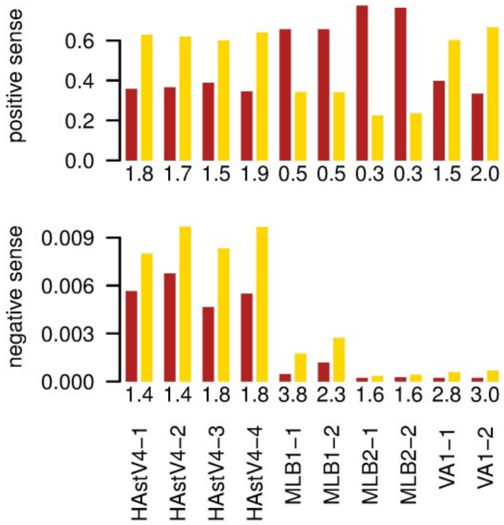


**C**

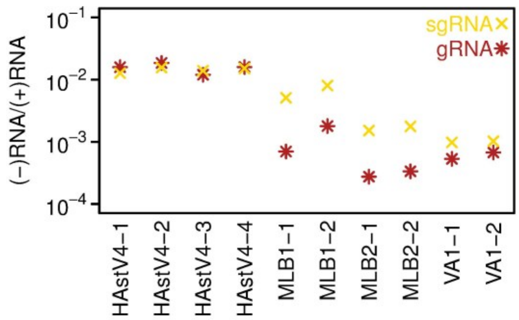

**Supplementary Figure 12. Total coverage of vRNA(+) and vRNA(−).** Caco-2 cells were infected with HAstV4 at MOI 5 and harvested at 24 hpi in quadruplicate. Fragments were mapped to vRNA(+) or vRNA(−), and total coverage summed. The y-axis scale is arbitrary but vRNA(−) coverage is scaled relative to vRNA(+) coverage by the indicated factor to aid visualization.

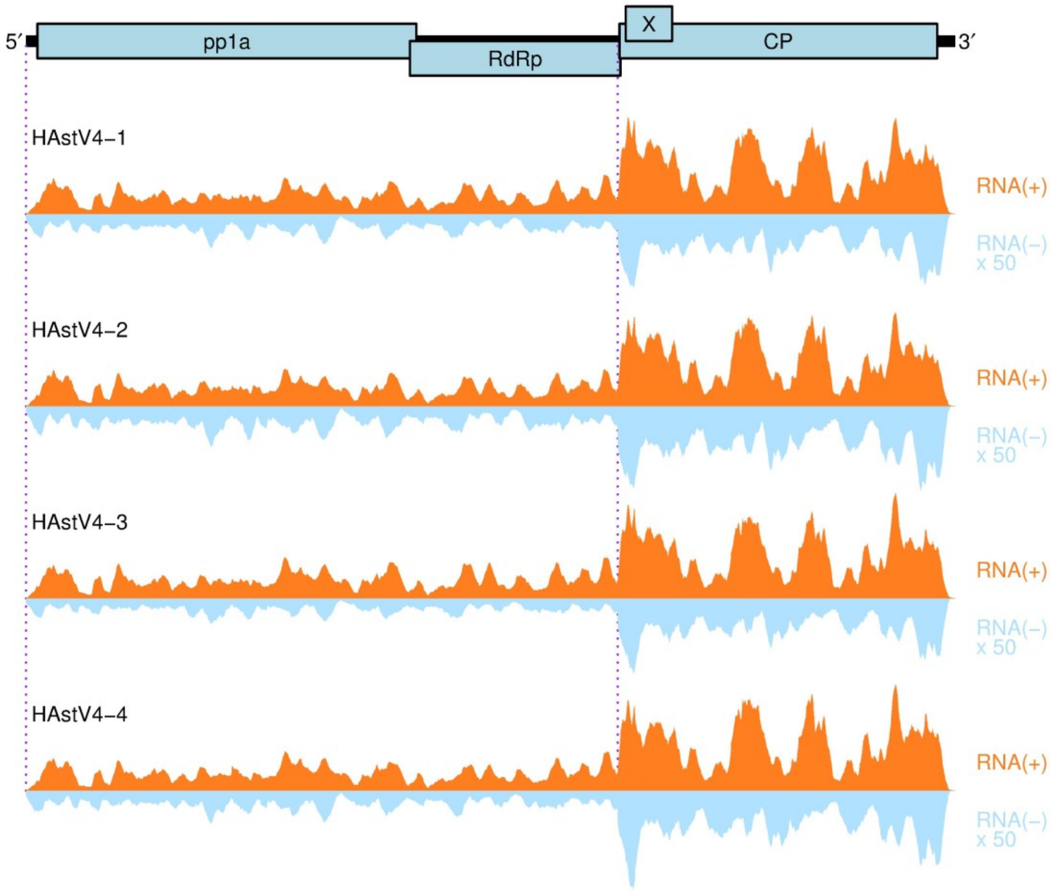


**Supplementary Figure 13. Total coverage of vRNA(+) and vRNA(−).** Huh7.5.1 cells were infected with MLB1 astrovirus at MOI 5 and harvested at 24 hpi in duplicate. Fragments were mapped to vRNA(+) or vRNA(−), and total coverage summed. The y-axis scale is arbitrary but vRNA(−) coverage is scaled relative to vRNA(+) coverage by the indicated factor to aid visualization.

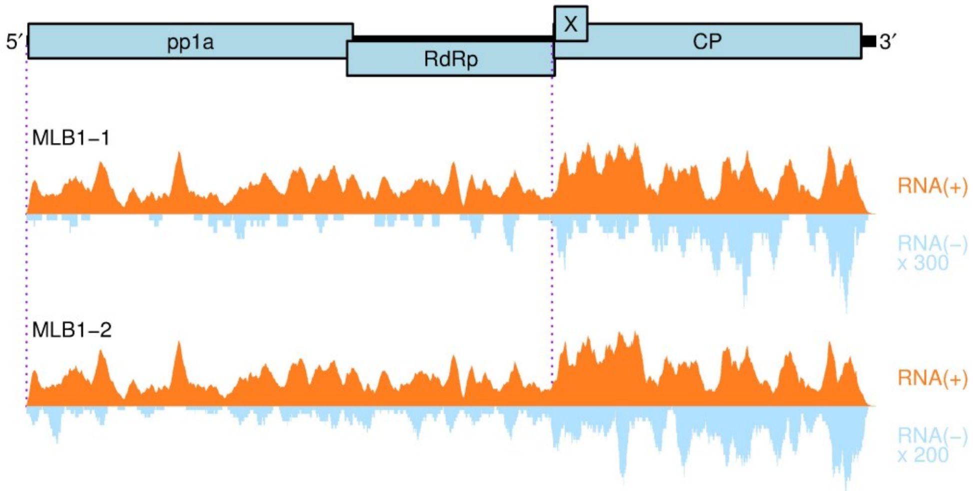

**Supplementary Figure 14. Total coverage of vRNA(+) and vRNA(−).** Huh7.5.1 cells were infected with MLB2 astrovirus at MOI 5 and harvested at 24 hpi in duplicate. Fragments were mapped to vRNA(+) or vRNA(−), and total coverage summed. The y-axis scale is arbitrary but vRNA(−) coverage is scaled relative to vRNA(+) coverage by the indicated factor to aid visualization.

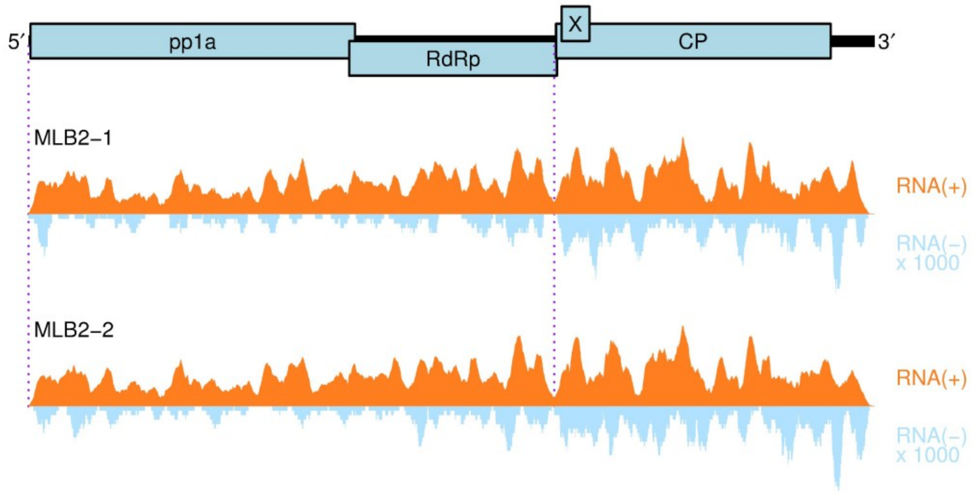


**Supplementary Figure 15. Total coverage of vRNA(+) and vRNA(−).** Caco-2 cells were infected with VA1 astrovirus at MOI 5 and harvested at 24 hpi in duplicate. Fragments were mapped to vRNA(+) or vRNA(−), and total coverage summed. The y-axis scale is arbitrary but vRNA(−) coverage is scaled relative to vRNA(+) coverage by the indicated factor to aid visualization.

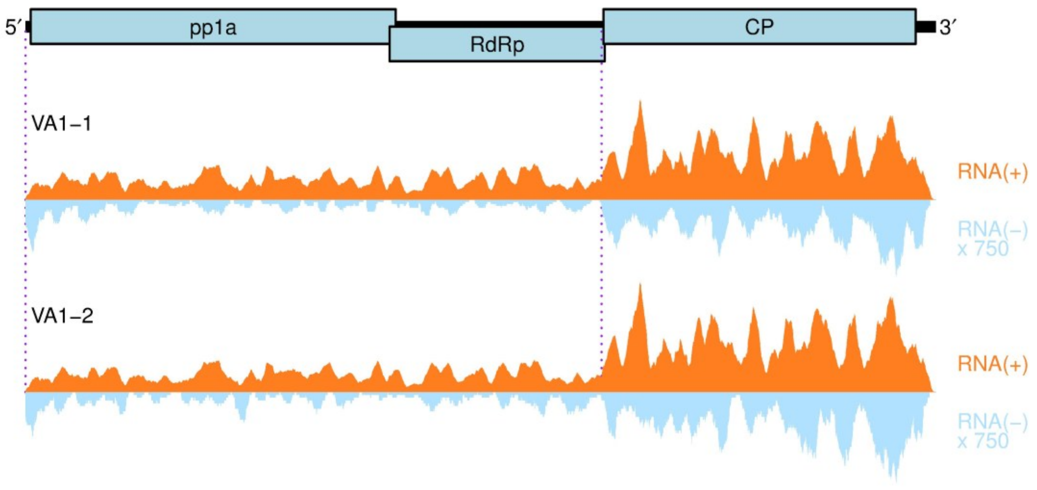

**Supplementary Figure 16. Histograms showing positions of 5′ ends of vRNA(+) fragments.** Caco-2 cells were infected with HAstV4 at MOI 5 and harvested at 24 hpi in quadruplicate. Counts are normalized to fragments per million fragments mapped to vRNA(+) or host mRNA(+) (RPM). Histograms show 5′ ends of positive-sense fragments.

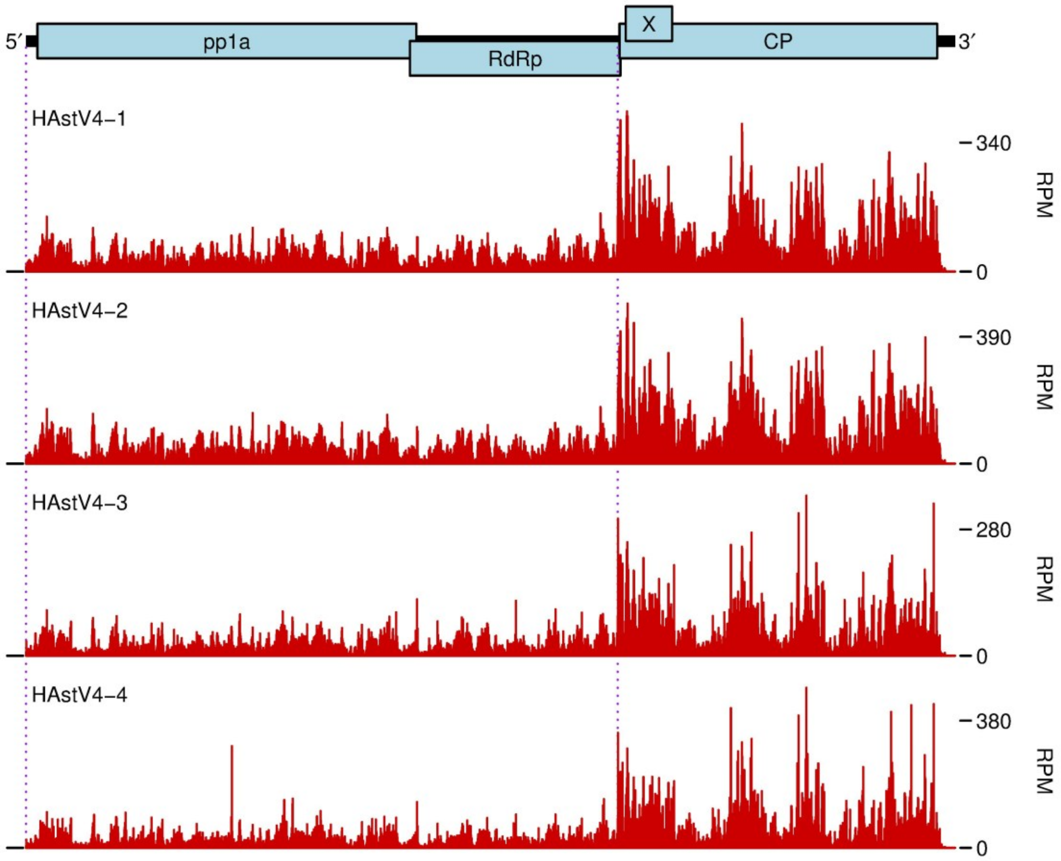

**Supplementary Figure 17. Histograms showing positions of 3′ ends of vRNA(−) fragments.** Caco-2 cells were infected with HAstV4 at MOI 5 and harvested at 24 hpi in quadruplicate. Counts are normalized to fragments per million fragments mapped to vRNA(+) or host mRNA(+) (RPM). Histograms show 3′ ends of negative-sense fragments, corresponding to 5′ ends of the positive-sense reverse complements of the fragments.

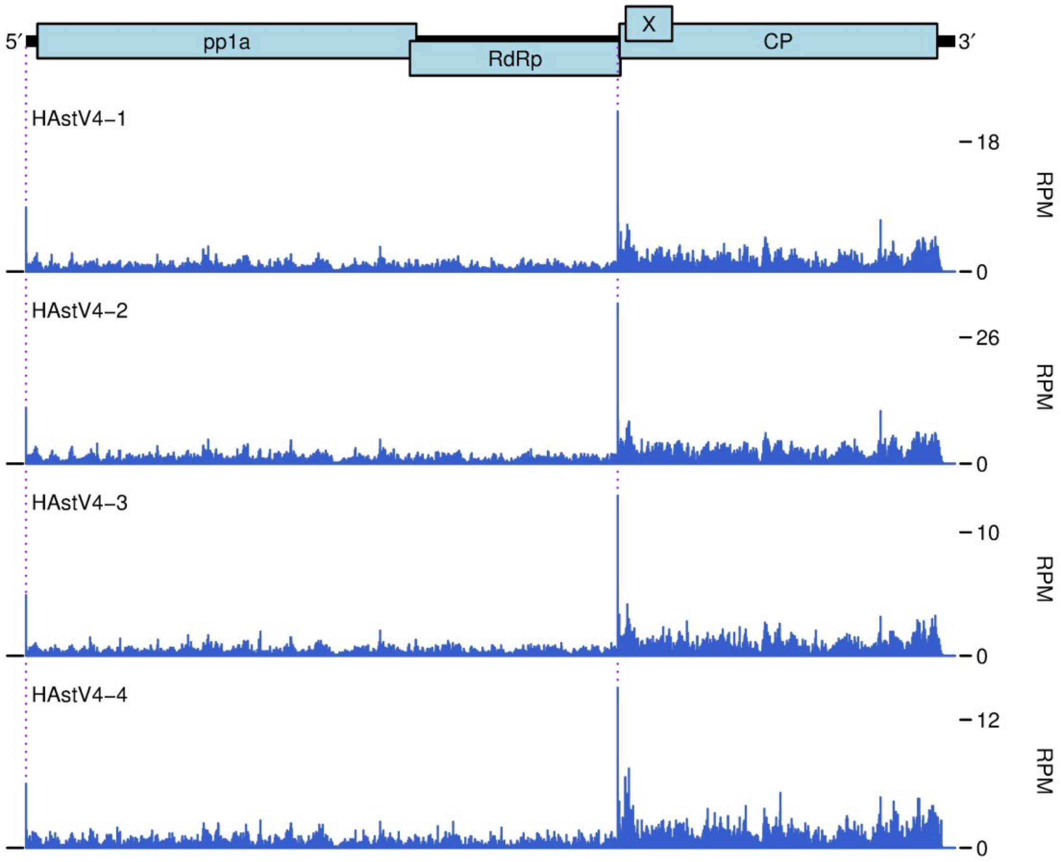

**Supplementary Figure 18. Histograms showing positions of 5′ ends of vRNA(+) fragments and 3′ ends of vRNA(−) fragments.** Huh7.5.1 cells were infected with MLB1 astrovirus at MOI 5 and harvested at 24 hpi in duplicate. Counts are normalized to fragments per million fragments mapped to vRNA(+) or host mRNA(+) (RPM). Histograms show 5′ ends of positive-sense fragments (red, upper plots) and 3′ ends of negative-sense fragments, corresponding to 5′ ends of the positive-sense reverse complements of the fragments (blue, lower plots).

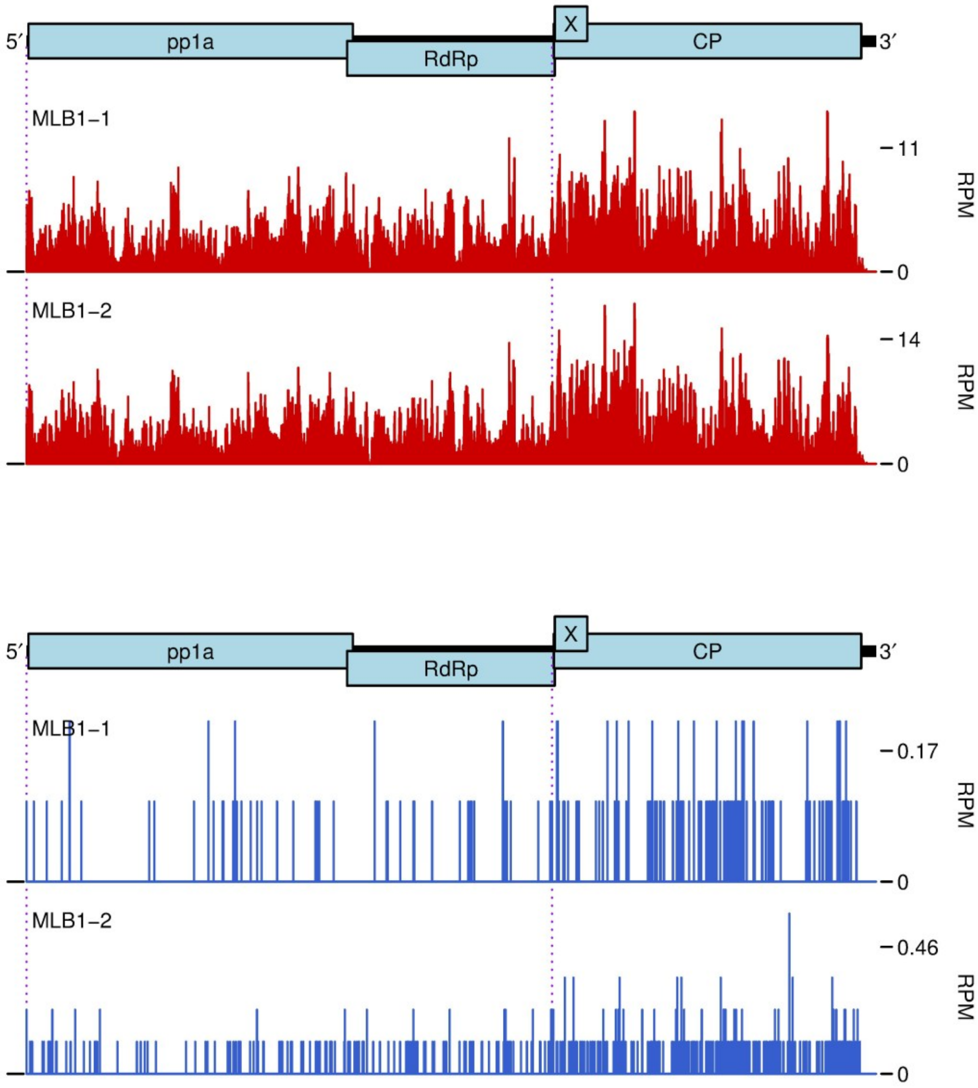

**Supplementary Figure 19. Histograms showing positions of 5′ ends of vRNA(+) fragments and 3′ ends of vRNA(−) fragments.** Huh7.5.1 cells were infected with MLB2 astrovirus at MOI 5 and harvested at 24 hpi in duplicate. Counts are normalized to fragments per million fragments mapped to vRNA(+) or host mRNA(+) (RPM). Histograms show 5′ ends of positive-sense fragments (red, upper plots) and 3′ ends of negative-sense fragments, corresponding to 5′ ends of the positive-sense reverse complements of the fragments (blue, lower plots).

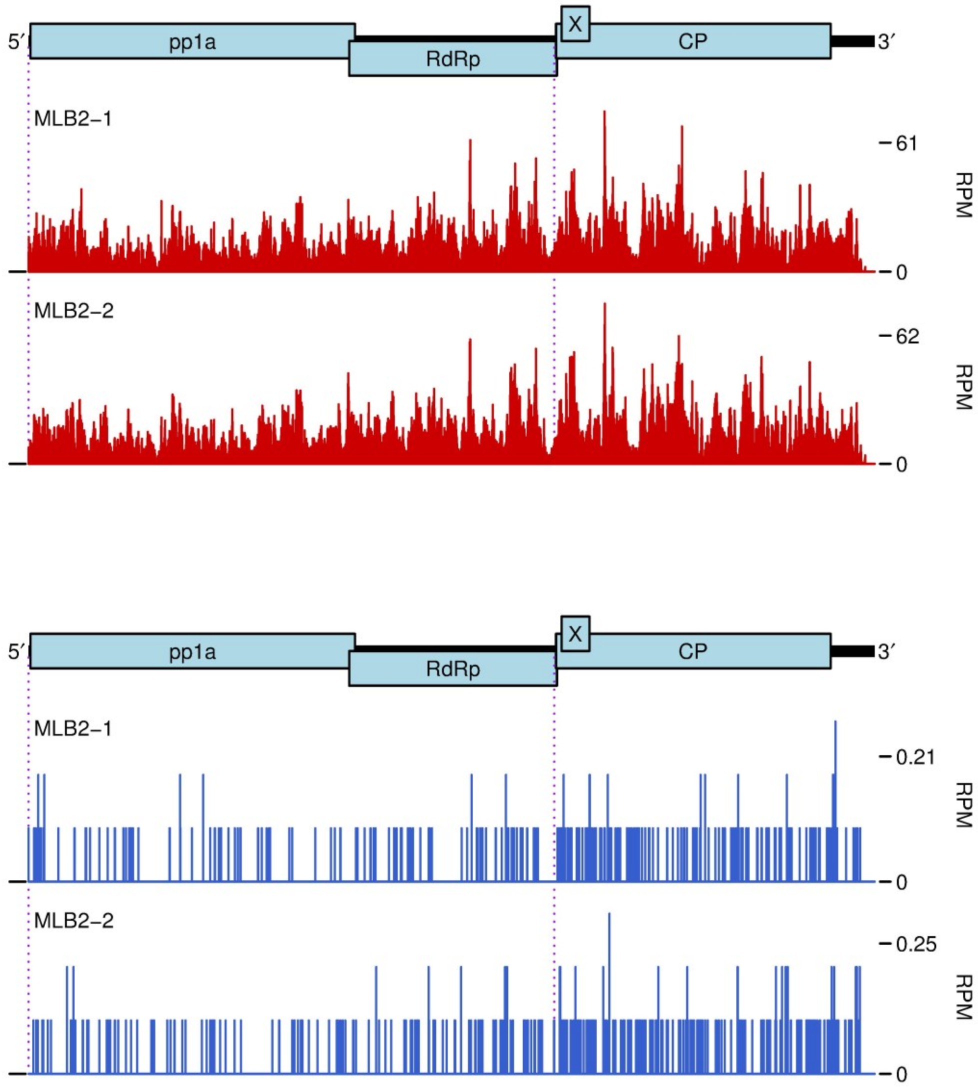

**Supplementary Figure 20. Histograms showing positions of 5′ ends of vRNA(+) fragments and 3′ ends of vRNA(−) fragments.** Caco-2 cells were infected with VA1 astrovirus at MOI 5 and harvested at 24 hpi in duplicate. Counts are normalized to fragments per million fragments mapped to vRNA(+) or host mRNA(+) (RPM). Histograms show 5′ ends of positive-sense fragments (red, upper plots) and 3′ ends of negative-sense fragments, corresponding to 5′ ends of the positive-sense reverse complements of the fragments (blue, lower plots).

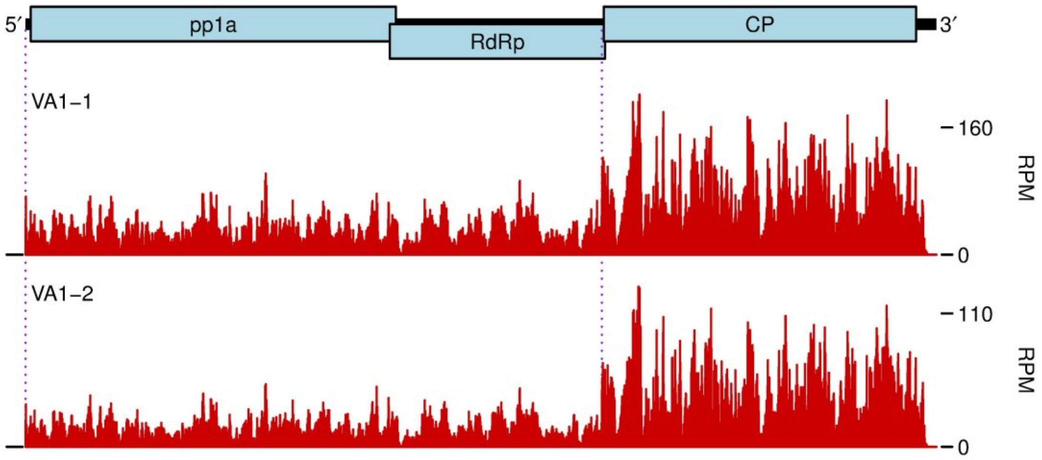


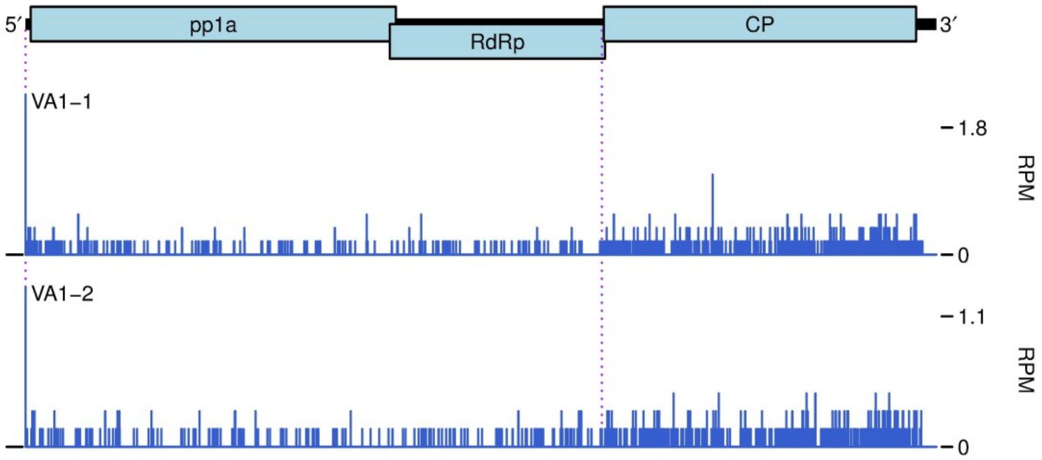

**Supplementary Figure 21. Read-length distributions of RNA-seq libraries.** Length distribution of virus-mapping fragments, after σ-clipping, for the first (**A**), second (**B**) and third (**C**) high throughput sequencing datasets.

**A**

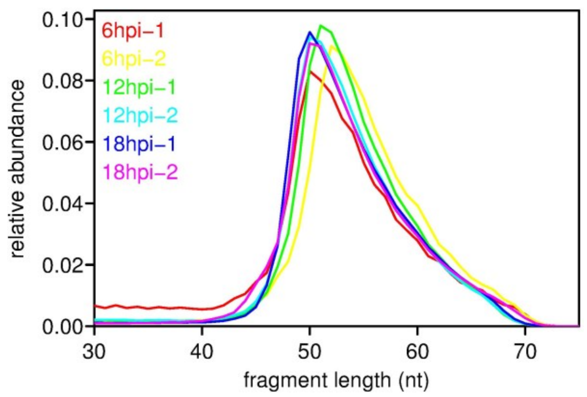


**B**

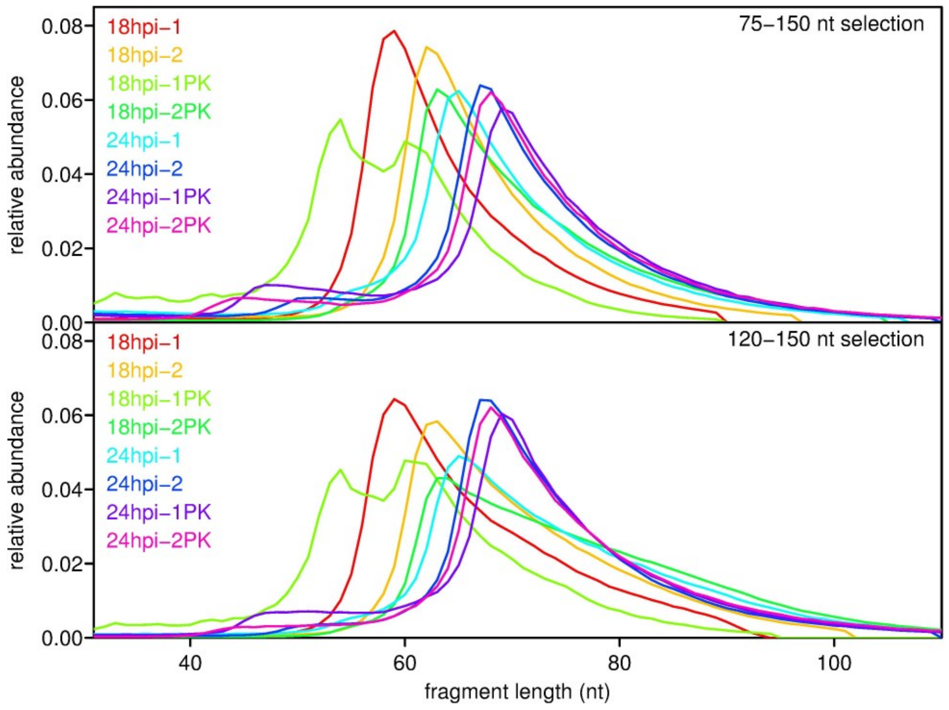


**C**

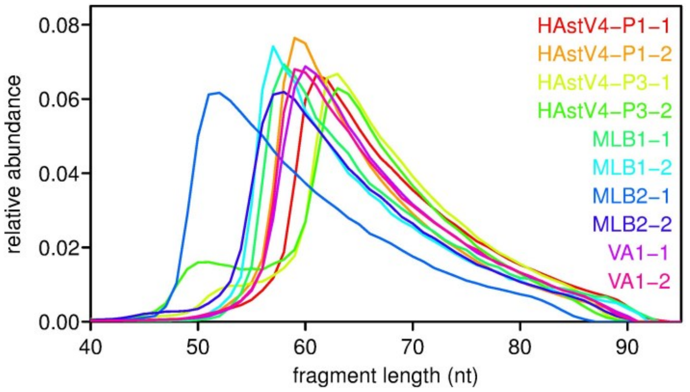

**Supplementary Table 1.** Host and virus read counts for the first high throughput sequencing dataset. Virus = HAstV1. Single-end sequencing. Total read counts are for the raw reads before any quality control or mapping.

| Timepoint | Repeat | Total reads | Host mRNA | vRNA(+) | vRNA(−) |
|---|---|---|---|---|---|
| 6 hpi | 1 | 35391199 | 5831197 | 52486 | 289 |
| 6 hpi | 2 | 41581106 | 6484718 | 67325 | 211 |
| 12 hpi | 1 | 15457347 | 3007362 | 630512 | 421 |
| 12 hpi | 2 | 11749063 | 2098351 | 469602 | 992 |
| 18 hpi | 1 | 16531373 | 2687283 | 1906618 | 11192 |
| 18 hpi | 2 | 24027747 | 3588304 | 2495754 | 17932 |

**Supplementary Table 2.** Host and virus read counts for the second high throughput sequencing dataset. Virus = HAstV1. Paired-end sequencing. Total read counts are the number of raw read pairs before any quality control or mapping. vRNA and mRNA read counts are for deduplicated and mapped read pairs.

| Timepoint | Repeat | Gel slice (nt) | Prot. K | Total read pairs | Host mRNA | vRNA(+) | vRNA(−) |
|---|---|---|---|---|---|---|---|
| 18 hpi | 1 | 75–150 | − | 12710966 | 1457826 | 2078280 | 11671 |
| 18 hpi | 2 | 75–150 | − | 9058305 | 1045556 | 1304349 | 1921 |
| 18 hpi | 1 | 75–150 | + | 10050648 | 702968 | 913427 | 22786 |
| 18 hpi | 2 | 75–150 | + | 6608003 | 731704 | 1946024 | 22196 |
| 24 hpi | 1 | 75–150 | − | 8946865 | 1171613 | 1401184 | 1655 |
| 24 hpi | 2 | 75–150 | − | 8280228 | 1053373 | 1215815 | 1726 |
| 24 hpi | 1 | 75–150 | + | 8712935 | 522112 | 722780 | 9717 |
| 24 hpi | 2 | 75–150 | + | 7151543 | 288727 | 470590 | 5297 |
| 18 hpi | 1 | 120–150 | − | 11103758 | 1480799 | 2097240 | 11353 |
| 18 hpi | 2 | 120–150 | − | 11191634 | 1682797 | 1998959 | 2908 |
| 18 hpi | 1 | 120–150 | + | 10759045 | 1008499 | 1259080 | 35844 |
| 18 hpi | 2 | 120–150 | + | 8627940 | 1024741 | 2685722 | 31178 |
| 24 hpi | 1 | 120–150 | − | 5862709 | 963411 | 1161242 | 1264 |
| 24 hpi | 2 | 120–150 | − | 7604984 | 1335713 | 1400545 | 1985 |
| 24 hpi | 1 | 120–150 | + | 6892147 | 509176 | 662140 | 8984 |
| 24 hpi | 2 | 120–150 | + | 11390288 | 662353 | 998886 | 11677 |

**Supplementary Table 3.** Host and virus read counts for the third high throughput sequencing dataset. Paired-end sequencing. Total read counts are the number of raw read pairs before any quality control or mapping. vRNA and mRNA read counts are for mapped read pairs.

| Virus | Timepoint | Repeat | Total reads | Host mRNA | vRNA(+) | vRNA(−) |
|---|---|---|---|---|---|---|
| HAstV4 | 24 hpi | 1 | 54319472 | 10058210 | 4597902 | 68077 |
| HAstV4 | 24 hpi | 2 | 29725722 | 4651497 | 2858424 | 50542 |
| HAstV4 | 24 hpi | 3 | 52589995 | 8809744 | 1994507 | 25517 |
| HAstV4 | 24 hpi | 4 | 36124894 | 5940483 | 1459639 | 23026 |
| MLB1 | 24 hpi | 1 | 47814255 | 9478754 | 140274 | 199 |
| MLB1 | 24 hpi | 2 | 46267125 | 8458245 | 158913 | 454 |
| MLB2 | 24 hpi | 1 | 66504586 | 10482777 | 690842 | 277 |
| MLB2 | 24 hpi | 2 | 71856494 | 9091993 | 673832 | 331 |
| VA1 | 24 hpi | 1 | 31106709 | 4184063 | 1109417 | 772 |
| VA1 | 24 hpi | 2 | 28990966 | 5872869 | 797653 | 659 |
